# A single vesicle fluorescence microscopy platform to quantify phospholipid scrambling

**DOI:** 10.1101/2025.04.17.649308

**Authors:** Sarina Veit, Grace I. Dearden, Kartikeya M. Menon, Faria Noor, Indu Menon, Takefumi Morizumi, Oliver P. Ernst, Anant K. Menon, Thomas Günther Pomorski

## Abstract

Scramblases play important roles in physiology by translocating phospholipids bidirectionally across cell membranes. For example, scrambling facilitated by dimers of the Voltage-Dependent Anion Channel 1 (VDAC1) enables endoplasmic reticulum-derived phospholipids to cross the outer membrane to enter mitochondria. Precise quantification of lipid scrambling, while critical for mechanistic understanding, cannot be obtained from ensemble averaged measurements of reconstituted scramblases. Here, we describe a microscopy platform for high-throughput imaging of single vesicles reconstituted with fluorescently labeled phospholipids and heterogeneously crosslinked VDAC1 dimers. For each vesicle, we quantify size, protein occupancy and scrambling rate. Notably, we find that individual VDAC1 dimers have different activities, ranging from <100 to >10,000 lipids per second. This kinetic heterogeneity, masked in ensemble measurements, reveals that only some dimer interfaces are capable of promoting rapid scrambling, as suggested by molecular dynamics simulations. We extend our analyses to opsin, a monomeric G protein-coupled receptor scramblase, thereby demonstrating the versatility of our platform for quantifying transbilayer lipid transport and exploring its regulation.

## Introduction

Scramblases facilitate the bidirectional translocation of polar lipids across cell membranes, a fundamental process that supports many aspects of cellular life. In metazoan cells, lipid scrambling is required for the exposure of the signaling molecule phosphatidylserine (PS) at the cell surface during apoptosis or upon cell activation, expansion of the endoplasmic reticulum (ER) and autophagosomal membranes, lipoprotein assembly, and all forms of protein glycosylation in the ER ^1–7^. Members of the G protein-coupled receptor (GPCR) ^8^ and TMEM16 ^9,10^ families were the first scramblases to be identified at the molecular level based on the demonstration of their scrambling activity after purification and reconstitution into unilamellar liposomes.

We recently reported that the mitochondrial β-barrel protein VDAC1 is a scramblase that provides the primary mechanism by which phospholipids, arriving at the mitochondrial surface from the ER via bridge-like transfer proteins ^11^, cross the outer membrane to enter mitochondria ^12^. Phospholipid import is needed to grow the double membrane system of mitochondria and provide precursors for the biosynthesis of cardiolipin, a tetra-acylated lipid that is necessary for mitochondrial function, as well as phosphatidylethanolamine (PE) which is needed in mitochondria but also distributed to other cellular membranes ^13,14^. Molecular dynamics simulations indicate that lipid scrambling is facilitated by VDAC1 dimers and occurs efficiently only at a specific dimer interface among several possible dimer configurations ^12^. To test this experimentally, we used an amine-reactive crosslinker to generate a mixture of VDAC1 dimers, including a subset with a suitable interface for lipid scrambling. When reconstituted into liposomes, ensemble average measurements revealed that the crosslinked (CL) VDAC1 preparation scrambled lipids, whereas the non-crosslinked (NCL) protein was inactive unless reconstituted at a sufficiently high protein-to-phospholipid ratio to allow spontaneous dimer formation ^12^. Given that VDAC1 is the most abundant protein in the mitochondrial outer membrane, with a density of >1,000 molecules per μm^2^ of surface ^15^, it is likely that scrambling-active native dimer interfaces result from molecular crowding.

Despite advances in the methodology to study lipid scramblases their functional and mechanistic characterization remains a challenge. Current approaches typically involve reconstituting purified scramblase proteins into large unilamellar liposomes (LUVs, approximately 200 nm in diameter) with defined bulk lipid compositions, followed by ensemble-averaged biophysical or biochemical studies. However, a fundamental limitation of ensemble measurements is the inherent heterogeneity of liposome preparations. Variability in size, lamellarity, membrane integrity, lipid composition, protein copy number, and protein orientation ^16–18^ complicates ensemble data interpretation by masking the dynamics of individual proteins. Importantly, such measurements also yield only average transport kinetics limiting the resolution and specificity of the analysis.

A single report in the literature describes an elegant attempt to overcome these challenges by analyzing the scramblase activity of Ca^2+^-activated TMEM16F in a microarray format, wherein purified protein is reconstituted into bespoke fabricated microchambers with an asymmetric bilayer consisting of a unique inner leaflet and an outer leaflet common to the entire microarray ^19^. Activation of the scramblase results in lipid transfer from the outer leaflet to the inner leaflet of the microchamber, which is tracked using fluorescent lipid reporters. Notably, these experiments yielded a unitary rate for TMEM16F-mediated scrambling, but the technical complexity of the set-up and demanding experimental workflow limit its practicality to specialized applications. Furthermore, the requirement for an activator to initiate scrambling precludes the use of this system for measuring constitutively active scramblases.

We now report a readily implementable, high-throughput single-vesicle assay to study scramblase activity, using dimeric human VDAC1 as an exemplar of scramblase proteins. We reconstituted vesicles with a fluorescent phospholipid reporter and fluorescently labeled, chemically crosslinked VDAC1 dimers, deposited the vesicles on a passivated glass slide and used total internal reflection fluorescence (TIRF) microscopy to resolve individual vesicles (Fig. 1, Extended Data Fig. 1). This approach enables a detailed characterization of the reconstituted proteoliposome ensemble, wherein we quantify vesicle size, protein occupancy, and lipid scrambling at the single-vesicle level using a novel adaptation of the common bovine serum albumin (BSA) back-extraction scramblase assay ^20,21^. By identifying vesicles containing a single VDAC1 dimer, we determined the number of lipids scrambled per dimer per second, i.e., the unitary scrambling rate. Notably, because our dimer preparation is heterogeneous, we found that the unitary rate ranged from <100 to >10,000 lipids per second, and that some dimers were inactive. This kinetic heterogeneity, which is masked in ensemble measurements, reveals that only a subset of dimer interfaces is capable of rapidly scrambling lipids, consistent with predictions from molecular dynamics simulations ^12^. To demonstrate the general applicability of our approach, we used the single vesicle platform to measure the scramblase activity of the GPCR opsin, which has been proposed to mediate lipid scrambling as a monomer ^22–24^. We show conclusively that monomeric opsin indeed scrambles phospholipids and that its unitary rate is much greater than that of optimal VDAC1 dimers, exceeding 10,000 lipids per second in most cases. Our platform can be readily modified and extended to quantify other phospholipid scramblases and determine, for example, how their activity is regulated by membrane lipid composition ^17^. It can also be used to study ATP-driven lipid transporters, i.e. flippases and floppases ^25,26^, providing a powerful and customizable method for their functional characterization.

**Figure 1:**
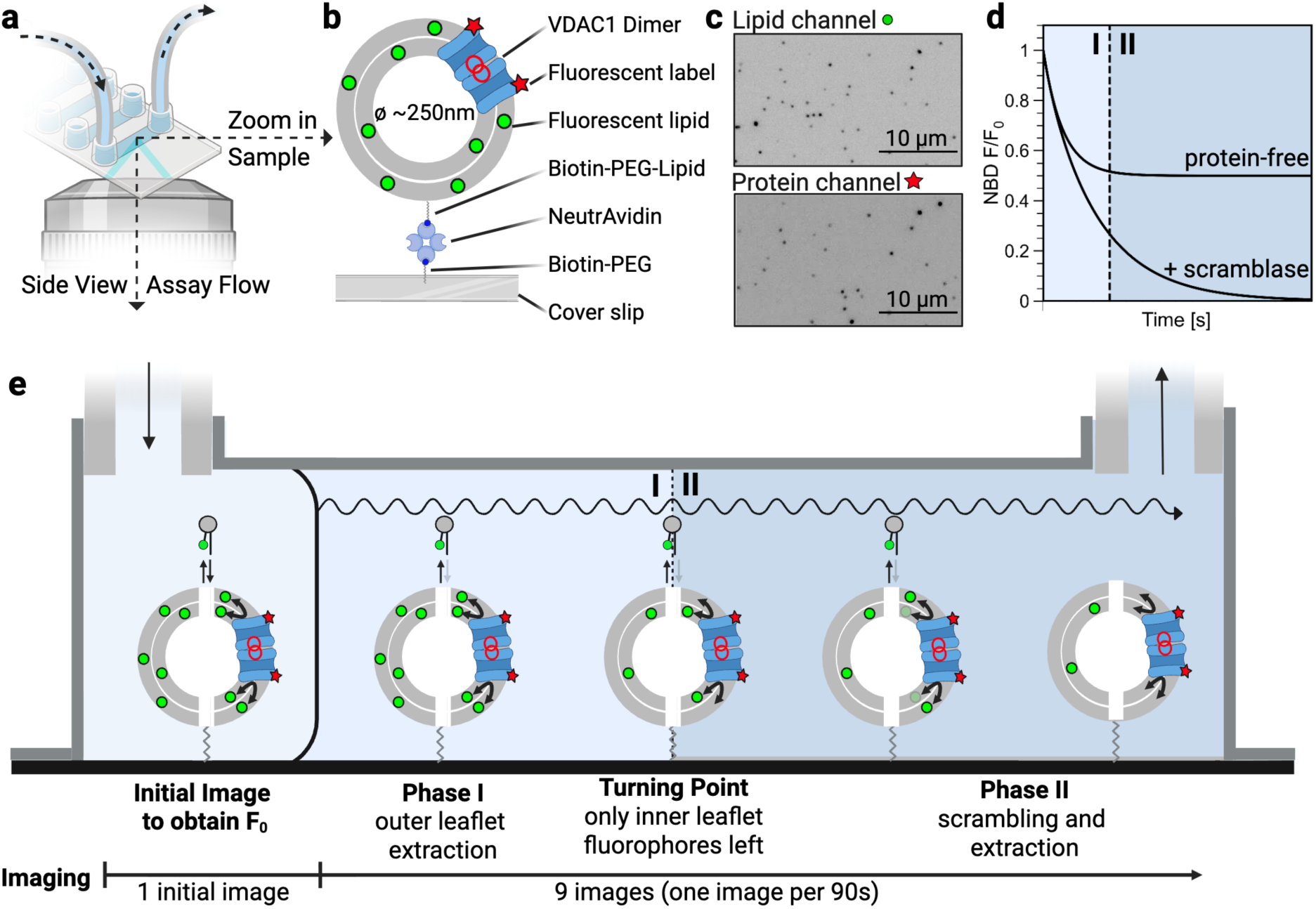
Single vesicle scramblase assay. **a**, Illustration of the sample chamber, with tubing for flushing, depicted in the context of a TIRF microscope objective lens (see also Extended Data Fig. 1). **b**, Illustration of a single large unilamellar vesicle (LUV), containing an Alexa Fluor 647-labeled crosslinked VDAC1 dimer (each protomer is modified by one fluorophore) and trace quantities of a fluorescent lipid (14:0-6:0 NBD-PC) and a biotinylated anchor lipid, immobilized on a polymer-passivated (PLL-g-PEG / PLL-g-PEG-biotin) glass cover slip via biotin-NeutrAvidin tethering. **c**, Representative TIRF images of immobilized vesicles visualized in the lipid channel (NBD) and protein channel (Alexa Fluor 647). The lipid signal serves as a read-out of scramblase activity, while the protein signal is used to determine the number of proteins per vesicle. **d**, Theoretical fluorescence traces for individual immobilized liposomes with (+scramblase) and without (protein-free) scramblase activity according to the protocol described in panel e. **e**, Schematic presentation of the assay steps. LUVs are introduced into the chamber where they become immobilized. An initial fluorescence image is captured to obtain the value of fluorescence (F_0_) at time = 0 s. Then, continuous flow of buffer (wavy line) through the imaging chamber removes 14:0-6:0 NBD-lipids that partition from the LUV outer leaflet into the buffer (phase I). At this juncture (turning point) the LUVs contain 14:0-6:0 NBD-lipids only in the inner leaflet. Next, in phase II of the sequence, these remaining 14:0-6:0 NBD-lipids are flipped to the outer leaflet in scramblase-containing LUVs where they become accessible to buffer extraction. Note that these phases partially overlap. The LUVs are depicted split down the middle to illustrate the outcome for those with a scramblase (right half) or without a scramblase (left half).

## RESULTS

### Single vesicle phospholipid scramblase assay

The fluorescent phospholipid analog 14:0-6:0 NBD phosphatidylcholine (NBD-PC) (Extended Data Fig. 2a) is commonly used as a reporter in ‘back-extraction’ phospholipid scramblase assays (Extended Data Fig. 2b) ^20,21,27^ because of its ability to desorb from membranes (rate constant ∼0.07 s^-1^, Extended Data Fig. 2c,d) into aqueous media where it has slight solubility (∼0.75 µM, Extended Data Fig. 2e). In such ‘back-extraction’ assays, large unilamellar vesicles (LUVs) composed typically of POPC and POPG (9:1, mol/mol) and containing trace amounts of NBD-PC (0.5 mole percentage of all lipids) are incubated with excess fatty acid-free albumin. NBD-PC molecules that desorb from the outer leaflet of the vesicle are captured within albumin’s hydrophobic binding pockets, leading to a decrease in fluorescence because the NBD fluorophore has a roughly 2-fold lower quantum efficiency when associated with albumin than when it is in the membrane (Extended Data Fig. 2b). For protein-free vesicles, where lipids do not scramble within the time-frame of the experiment, about 25% of the sample fluorescence is lost (Extended Data Fig. 2c) as inaccessible NBD-PC molecules in the inner leaflet of the vesicles are retained while those in the outer leaflet are captured by albumin. However, if a vesicle possesses a scramblase, NBD-PC molecules in the inner leaflet are translocated to the outer leaflet, ultimately allowing the entire pool to be captured. This results in about 50% fluorescence quenching (Extended Data Fig. 3f, ‘CL’ trace). Building on this well-characterized system, we developed a new method to visualize VDAC1-mediated phospholipid scrambling in single vesicles.

Our method requires LUVs reconstituted with the NBD-PC reporter, a biotinylated anchor lipid, and a fluorescently labeled scramblase (protein-free LUVs serve as a control). A suspension of the LUVs is introduced into a flow chamber (Fig. 1a, Extended Data Fig. 1), the floor of which is a passivated glass cover slip coated with NeutrAvidin (Fig. 1b). The LUVs become immobilized as their biotinylated anchor lipid attaches to the NeutrAvidin coating (Fig. 1b) and can then be imaged by total internal reflection fluorescence (TIRF) microscopy in both lipid and protein fluorescence channels (Fig. 1c). With LUVs immobilized on the cover slip, a small pool of NBD-PC is expected to be in the aqueous phase in equilibrium with the NBD-PC residing in the outer leaflet of the LUV membrane (Fig. 1e, corresponding to the scenario in ‘Initial Image’). On flowing buffer through the chamber, this aqueous pool of NBD-PC will be washed away, driving the progressive depletion of outer-leaflet NBD-PC as it continues to desorb into solution (Fig. 1d,e, ‘Phase I’). Thus, initially, all vesicles will lose their complement of outer leaflet NBD-PC, resulting in a ∼50% loss of fluorescence (Fig. 1d, ‘protein-free’). In LUVs with scramblase activity, inner leaflet NBD-PC will translocate outwards, desorb, and be washed away, resulting in complete loss of fluorescence (Fig. 1d,e, ‘Phase II’). If scrambling is slower than the rate of outer-leaflet depletion of NBD-PC, the two processes can be deconvoluted, yielding a scrambling rate constant. By identifying vesicles equipped with a single VDAC1 dimer and knowing the size of the vesicle and therefore the number of phospholipids it contains, we expect to be able to determine the unitary rate of VDAC1-mediated scrambling.

### Validation of the experimental platform

We validated our experimental platform in two ways. First, we confirmed that NBD-lipids are lost from the outer leaflet via desorption and buffer flow, resulting in the predicted loss of fluorescence from individual vesicles. To test this, we prepared LUVs labeled symmetrically with NBD-PC, or with head-labeled *N*-NBD-PE which cannot desorb readily as it has two long acyl chains which confine it to the membrane. We also prepared asymmetrically labeled LUVs with NBD-PC located only in the outer leaflet. After immobilizing the LUVs in the sample chamber, we initiated imaging and buffer flow (500 µl/min peristaltic pump flow rate, 24°C). *N*-NBD-PE labeled LUVs maintained a relatively stable fluorescence signal throughout the observation period (Fig. 2a), consistent with the lipid remaining associated with the LUVs. In contrast, LUVs asymmetrically labeled with NBD-PC showed a complete loss of fluorescence within the first 200 s (Fig. 2b). Finally, liposomes symmetrically labeled with NBD-PC, showed an initial rapid decrease in fluorescence (within the first 200 s), reaching a plateau at 49.5 ± 8.5 % (mean ± S.D., n=310) of the initial signal, corresponding to the non-extractable lipid pool in the inner leaflet (Fig. 2c). Mono-exponential fitting of the time-dependence of fluorescence loss from the NBD-PC-containing samples yielded a rate constant γ ∼0.0108 ± 0.0045 s^-1^ (mean ± S.D., n=394), characterizing the composite process of desorption of the lipid and its removal by buffer flow.

**Fig. 2.**
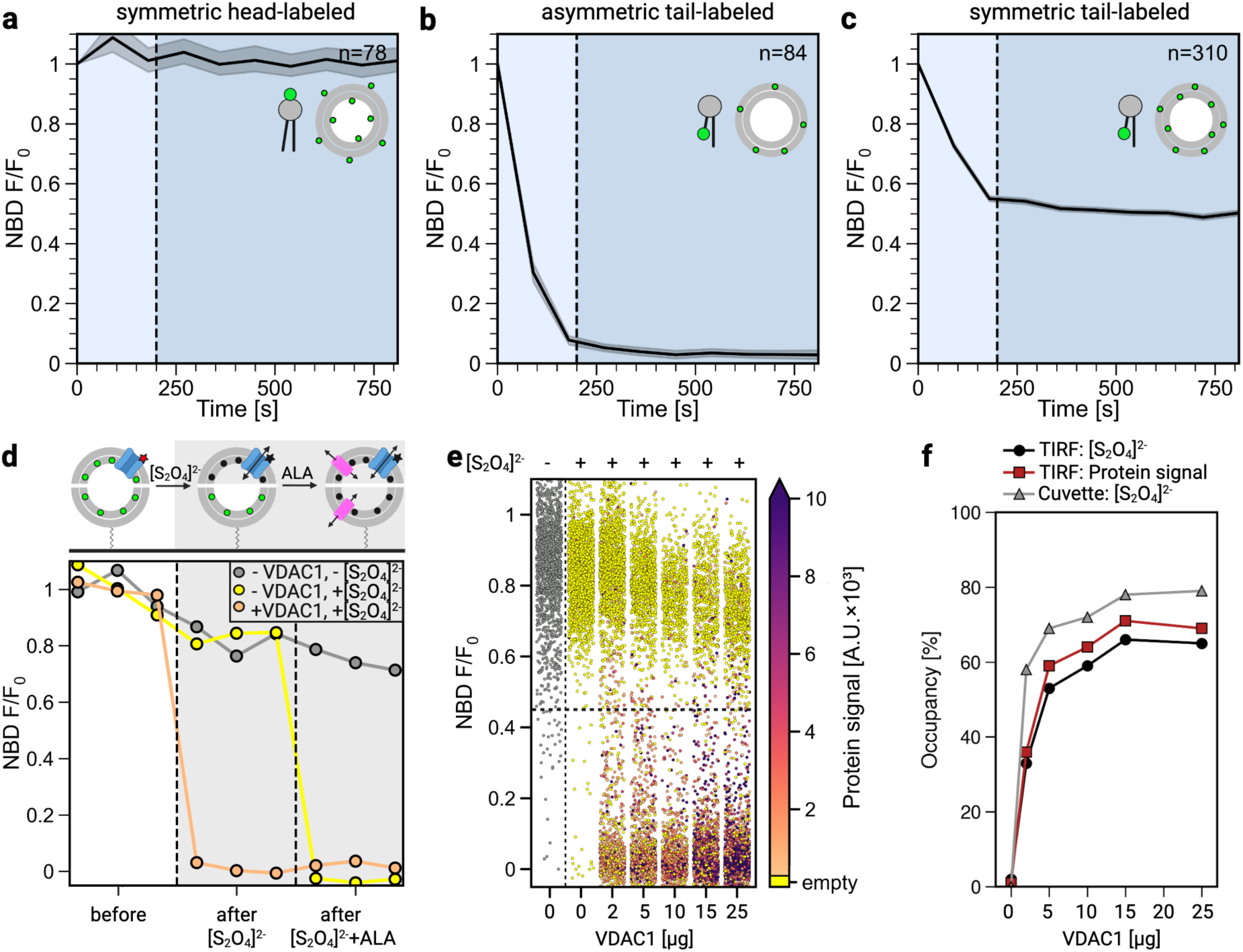
Validation of the single vesicle scramblase assay platform. **a-c**, LUVs labeled symmetrically or asymmetrically with the indicated NBD-lipids were immobilized as in Figure 1b,e. TIRF microscopy imaging was started simultaneously with buffer flow (t=0 min). For each vesicle, lipid fluorescence intensity was normalized to the corresponding initial intensity, and individual traces were combined into a plot showing the mean. The 95% confidence interval is indicated by the shaded error band and the number of averaged individual LUV traces is indicated at the top right of the graphs. **a**, Symmetrically-labeled LUVs with head-labeled *N*-NBD-PE in both leaflets. **b**, Asymmetrically-labeled LUVs with 14:0-6:0 NBD-PC solely in the outer leaflet. **c**, Symmetrically-labeled LUVs with 14:0-6:0 NBD-PC in both leaflets. **d**, Schematic representation of protocol (top panel) and exemplary data (bottom panel). Note that the LUVs in the top panel are depicted split through the middle to illustrate the outcome for those with (top half) or without (bottom half) a VDAC1 channel. 14:0-6:0 NBD-PC-containing proteoliposomes reconstituted with Alexa Fluor 647-labeled VDAC1 (non-crosslinked) were imaged by TIRF microscopy alongside protein-free liposomes to determine the efficiency of VDAC1 reconstitution. After vesicle immobilization, buffer flow was used to remove outer-leaflet NBD-PC molecules (the resulting vesicles are depicted at the start of the sequence in the top panel). This was followed by dithionite ([S_2_O_4_]^2-^) addition to bleach NBD-PC in the inner leaflet of VDAC1-containing vesicles (dithionite enters the vesicles via the VDAC1 channel). Alamethicin (ALA) was then added to permeabilize all vesicles, confirming complete bleaching. Exemplary fluorescence traces are shown for a VDAC1-containing proteoliposome (orange), a protein-free liposome (yellow), and a control protein-free liposome sample (grey) that was not treated with [S_2_O_4_]^2-^. The latter control (grey) indicates the extent of photobleaching through the imaging process. See also Extended Data Figure 4a,b. **e**, NBD fluorescence remaining after adding [S_2_O_4_]^2-^ to LUVs reconstituted with the indicated amounts non-crosslinked Alexa Fluor 647-labeled VDAC1, with the initial protein signal indicated by a colormap (scale on the right; the protein colormap corresponds to the initial Alexa Fluor 647 fluorescence of the vesicles as this fluorophore is also bleached on dithionite treatment). The control sample indicated in the first column (grey dots) was not subjected to [S_2_O_4_]^2-^ treatment. The dashed line indicates the 45% cut-off. **f**, Comparison of vesicle protein occupancy determined by single vesicle assay using [S_2_O_4_]^2-^ (black), single vesicle protein signal (red) and ensemble [S_2_O_4_]^2-^ assay (grey). The x-axis represents the amount of VDAC1 used per reconstitution.

The value of the rate constant γ that we measured is smaller than the desorption rate constant determined in ensemble assays (Extended Data Fig. 2d). To investigate this, we tested the effect of changing the buffer flow rate and temperature and found that an increase in either of these parameters results in an increase in γ (Extended Data Fig. 2f,g). However, as higher flow rates affect microscope stage stability and image focus, we picked a flow rate that was optimized to balance efficient lipid desorption with minimal mechanical disturbance. We also considered fluorescent lipid reporters other than NBD-PC. As phospholipid scramblases are generally unspecific ^10,12,28^ we reasoned that it should be possible to choose between several short-chain NBD lipids, to identify those with the highest γ value in order to have the best time resolution for the assay. As shown in Extended Data Fig. 2h, increasing the length of the 1-acyl chain of various NBD lipids from C14 to C16 predictably decreased γ. However, the extraction rates of 14:0-6:0 NBD-PE and C6-NBD-sphingomyelin were higher (Extended Data Fig. 2h); the higher γ value for the latter compared with that of the corresponding NBD-PC is likely because of the C3-hydroxyl in its sphingosine backbone which gives it greater polarity. These results validate the behavior of NBD-lipids, specifically our principal reporter NBD-PC, in the single vesicle assay system.

We next tested whether the single vesicle platform could recapitulate previously reported ensemble data on VDAC1 reconstitution ^12^. For this we needed to determine the fraction of vesicles functionalized with a VDAC1 channel and relate this information to the amount of protein used for reconstitution. Thus, we purified a single-cysteine variant of VDAC1 (Extended Data Fig. 3a,b) in which the cysteine residue (A170C) faces the aqueous pore of the β-barrel where it does not interfere with scramblase activity. We used Alexa Fluor 647-maleimide to label the A170C site with high efficiency (>97%, Extended Data Fig. 3c) and reconstituted the protein into LUVs containing NBD-PC. Under these conditions, VDAC1 functions solely as a channel as its scramblase activity requires dimerization, which can be achieved by chemical crosslinking (Extended Data Fig. 3e,f). We immobilized the VDAC1 proteoliposomes onto a TIRF microscope channel slide (Fig. 1a,b, Extended Data Fig. 1c) and flushed the chamber with buffer for 5 min to remove NBD-PC molecules from the outer leaflet of the vesicles. After stopping buffer flow, we carried out a staged protocol (detailed in Fig. 2d (schematic, top panel) and Extended Data Figure 4) as follows. First, we imaged the sample to determine the lipid and protein fluorescence of each LUV (’before’ image). We next added dithionite ([S_2_O_4_]^2^), a membrane impermeant reductant which reacts with NBD fluorophores to eliminate their fluorescence (Fig. 2d, Extended Data Fig. 3e, Extended Data Fig. 4) and measured the remaining fluorescence in each vesicle ^18^ (’after [S_2_O_4_]^2-^’ image) – we expected that protein-free vesicles in the sample would retain their fluorescence whereas vesicles with VDAC1 channels would become non-fluorescent. Finally, we added the [S_2_O_4_]^2-^-permissive pore-forming peptide alamethicin (ALA)(Fig. 2d, Extended Data Fig. 4) to expose all pools of NBD-PC to [S_2_O_4_]^2-^ and again measured the remaining fluorescence in each vesicle (’after ALA’ image).

On testing the protocol with protein-free vesicles in the absence of dithionite, we expected to see no change in fluorescence but in practice observed ∼8% loss of fluorescence on average (Extended Data Fig. 4a, ‘before’) with the remaining fluorescence being 91.6 ± 0.85 % (mean ± SEM, n=1,498) of the starting value (Fig. 2d, bottom panel, grey trace; Fig. 2e, first column). A similar pattern was seen when protein-free vesicles were treated with dithionite (Fig. 2d, bottom panel, yellow trace; Fig. 2e, second column, Extended Data Fig. 4b). In a very small subset of vesicles, we found that as much as ∼55% of the fluorescence was lost, likely because NBD-PC molecules in the outer leaflet of these vesicles had not been completely removed during the initial phase of the experiment (Fig. 2e, first column, Extended Data Fig. 4a). Accordingly, we chose 45% residual fluorescence as a cut-off (Extended Data Fig. 4a,b, dashed line indicates cut-off) to distinguish protein-free vesicles from those containing VDAC1 channels which permit [S_2_O_4_]^2-^ entry, and which consequently lose all fluorescence on dithionite treatment (Fig. 2e). We found that the latter vesicles had a signal >300 in the Alexa Fluor 647 channel, indicating that this threshold is a reliable estimate of the minimum fluorescence signal of a single Alexa Fluor 647-VDAC1 protein (see also later).

Reconstitution of VDAC1 into vesicles is proposed to follow Poisson statistics, with the fraction of vesicles containing one or more VDAC1 molecules being correlated with the amount of protein used. To test this, we prepared vesicles with different protein/phospholipid ratios and used Alexa Fluor 647 imaging to measure directly the percentage of vesicles containing at least one VDAC1, i.e., the percentage of vesicles with an Alexa Fluor 647 signal >300 units. We treated the immobilized samples with [S_2_O_4_]^2^, as described above, to determine the fraction of vesicles that lost their entire NBD fluorescence, indicating the presence of at least one functional VDAC1 channel. When graphed as a function of the amount of Alexa Fluor 647-VDAC1 used for reconstitution, both the direct measurement of Alexa 647 fluorescence intensity of vesicles (Fig. 2f, red squares) and the functional read-out obtained via [S_2_O_4_]^2-^-mediated bleaching (Fig. 2f, black circles) yielded identical profiles. We compared these single-vesicle approaches with results obtained from bulk assays where [S_2_O_4_]^2-^treatment was used to infer the fraction of vesicles containing VDAC1 channels (Fig. 2f, grey triangles; Extended Data Fig. 3e). Compared to the ensemble analysis, the immobilized liposomes showed a slightly lower protein occupancy (Fig. 2f) and an enrichment of smaller vesicles (Extended Data Figure 5e). This enrichment of small vesicles, also noted in previous studies ^18,29,30^, is likely due to their faster diffusion, which promotes more efficient immobilization. We conclude that our single-vesicle setup enables both direct quantification and functional assessment of protein content.

### Visualization of phospholipid scrambling by cross-linked VDAC1 in single vesicles

Under our standard reconstitution conditions, crosslinked VDAC1 (CL-VDAC1) exhibits scramblase activity in ensemble assays, whereas non-crosslinked VDAC1 (NCL-VDAC1) does not (Extended Data Fig. 3f) ^12^. To generate CL-VDAC1, we treated Alexa Fluor 647-labeled VDAC1 with EGS, an amine-reactive homo-bifunctional crosslinker, generating a sample with a significant proportion of crosslinked protein (Extended Data Fig. 3d). Crosslinking has no discernible effect on channel activity as determined by the dithionite permeation assay (Extended Data Fig. 3e). When reconstituted and assayed in bulk via the BSA-back-extraction assay, CL-VDAC1 displayed expected scramblase activity, whereas NCL-VDAC1 was inactive (Extended Data Fig. 3f). The CL-Alexa Fluor 647-VDAC1 preparations used in our experiments consist of a roughly equal mixture of crosslinked and non-crosslinked protein (Extended Data Fig. 3d). To refine our analysis and measure the rate of lipid scrambling catalyzed by a single VDAC1 dimer, we purified the crosslinked dimer fraction (henceforth *p*urified *c*ross*l*inked, pCL) by size-exclusion chromatography (Extended Data Fig. 5a) and verified its channel and scramblase activity in cuvette assays (Extended Data Fig. 5b,c).

To visualize scramblase activity in single vesicles, we reconstituted NBD-PC-vesicles with pCL-Alexa Fluor 647-VDAC1 and introduced these into a TIRF chamber, allowing 1 min for the vesicles to reach the NeutrAvidin-coated surface. This was followed by a 1-min pause to enable immobilization before acquiring an initial NBD fluorescence image (Fig. 1e, initial image). We used mock-reconstituted liposomes and NCL-Alexa Fluor 647-VDAC1 proteoliposomes as controls. Buffer flow was started immediately after acquiring the initial image and 9 additional images were captured at 90 second intervals, a frequency optimized to minimize photobleaching effects, which accounted for a 1.0 ± 13.4 % (n=63) loss in signal at the end of the image acquisition sequence, as determined using vesicles containing non-extractable head-labeled *N*-NBD-PE (Fig. 2a). After NBD imaging, Alexa Fluor 647 fluorescence in each vesicle was quantified to estimate VDAC1 content based on initial protein signal as well as Alexa Fluor 647 photobleaching step analysis.

For protein-free liposomes and NCL-Alexa Fluor 647-VDAC1 proteoliposomes, decay of NBD fluorescence stabilized in ∼100 s, reaching a plateau (determined at 900 s) at 47 ± 10 % (n=254) and 42 ± 9% (n=283) of the initial fluorescence signal, respectively (Fig. 3a,b). This indicates the expected loss of only outer-leaflet NBD-PC, consistent with the absence of scrambling in these control samples. In contrast, proteoliposomes reconstituted with pCL-Alexa Fluor 647-VDAC1 showed deep fluorescence reduction, indicating that inner-leaflet NBD-PC was translocated to the outer leaflet and subsequently lost from the vesicle. The greatest extents of reduction within the 900 second assay window corresponded to vesicles with high protein content (Fig. 3c). For simplified presentation, we grouped fluorescence traces by VDAC1 content based on Alexa Fluor 647 fluorescence, with category averages displayed in Figure 3d-f. We conclude that our experimental platform enables clear visualization of the protein-dependent scramblase activity of pCL-Alexa Fluor 647-VDAC1 in single vesicles.

**Figure 3.**
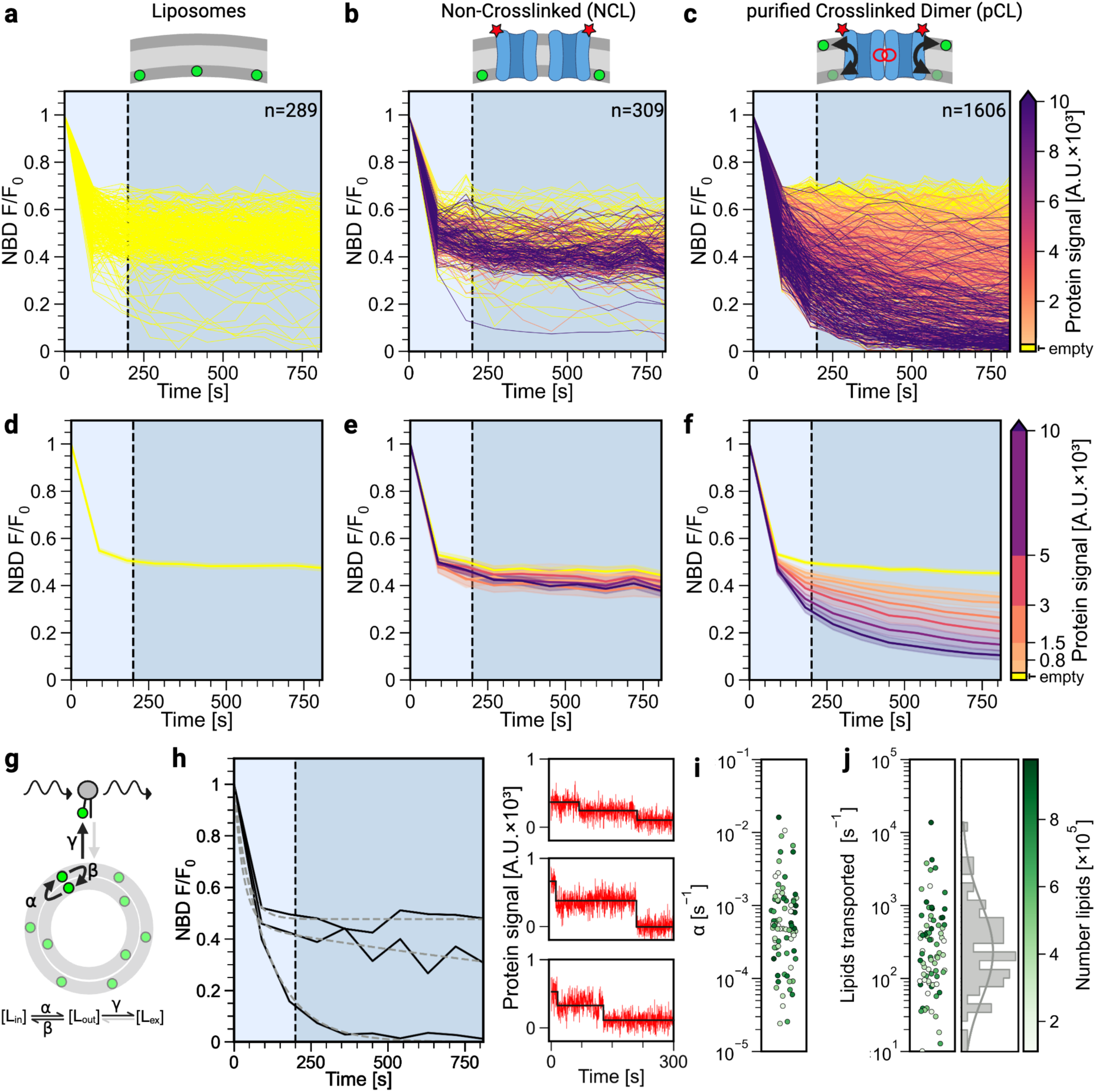
Single vesicle analysis of VDAC1-mediated scramblase activity. 14:0-6:0 NBD-PC proteoliposomes containing either non-crosslinked (NCL) or purified crosslinked (pCL) Alexa Fluor 647-labeled VDAC1 dimers, as well as protein-free NBD-PC liposomes were immobilized and subjected to TIRF microscopy as illustrated in Figure 1e. Buffer flow was initiated after acquiring an initial image at t=0 min, and subsequent images were taken every 90 s. Panels show time-dependence of the NBD signal for (**a, d**) protein-free liposomes, (**b, e**) NCL-VDAC1 proteoliposomes, and (**c, f**) pCL-VDAC1 dimer proteoliposomes. For each individual liposome, NBD fluorescence intensity is shown as a normalized line plot relative to the initial fluorescence (F_0_). Protein occupancy in each liposome is color-coded based on Alexa Fluor 647 signal intensity, as indicated by the colormap on the right of panels **c** and **f**. **d - f**, Fluorescence traces from vesicles with similar protein occupancy were grouped and plotted in increasing order of protein content, with the protein signal and intensity grouping indicated by the colormap on the right. The shaded error band indicates the 95% confidence interval. **g**, Schematic representation of the three-compartment model (Supplementary Information) used for data fitting, illustrating the inner leaflet (L_in_), outer leaflet (L_out_) and extracted pool (L_ex_), along with the respective rate constants. Fits were done by setting the scrambling rate constants α=β and using constraints specified in Model 1 (Extended Data Table 1). **h**, Representative NBD-fluorescence intensity traces (solid lines) fitted using the three-compartment model (dashed lines) for three single vesicles exhibiting two-step Alexa Fluor 647 photobleaching events (panels on the right), indicative of the presence of a single pCL-VDAC1 dimer in the vesicles. **i**, Scrambling rate constant α for active vesicles with a single pCL-VDAC1 dimer. Symbols are color-coded to indicate number of lipids per vesicle according to the color map in panel j. **j**, Number of transported lipids per second for individual vesicles with a single pCL-VDAC1 dimer, color-coded to indicate number of lipids per vesicle according to the color map on the right with histogram illustrating the distribution of values. In (**i, j**) the colormap represents the total number of lipids for each vesicle, deduced from initial NBD fluorescence intensity and dynamic light scattering data (see Extended Data Fig. 5e).

### Unitary rate of scrambling by a VDAC1 dimer

To derive the scrambling rate from the traces shown in Figure 3c, we built a three-compartment kinetic model (Fig. 3g, Supplementary Information) comprising two pools of fluorescent NBD-PC (L_in_ and L_out_, representing lipids in the inner and outer leaflet of a single vesicle, respectively) and a third pool (L_ex_) representing NBD-PC molecules that desorb from the outer leaflet and are removed by buffer flow. Notably, the L_ex_ pool does not contribute to the vesicle fluorescence. To avoid overfitting as each trace has only 10 data points, we assumed that scramblase-mediated exchange between the L_in_ and L_out_ pools occurs at an equal rate in both directions (rate constants α = β), with an initial lipid distribution of 0.4 < L_out_/(L_in_ + L_out_) < 0.6. We used Akaike and Bayesian Information Criteria to evaluate three fitting models with different combinations of constraints for the extraction rate constant γ and the scrambling rate constant α, and identified one model for our analyses (Extended Data Table 1).

We obtained high-quality fits (r² > 0.8). However, although protein occupancy increased with vesicle diameter (Extended Data Fig. 5f), the data revealed significant heterogeneity in scrambling rates with only a weak correlation between protein content and scrambling efficiency (Extended Data Fig. 5g). We attribute this heterogeneity to differences in dimer interfaces generated during EGS crosslinking. Thus, using coarse-grained molecular dynamics simulations we previously predicted ^12^ that the scrambling rate is extremely sensitive to the nature of the interface – for example, the simulations indicated robust scrambling by an optimal VDAC1 dimer, while slight rotations of the individual protomers away from this optimal arrangement, generated interfaces with 5-fold reduced scramblase activity ^12^. Likewise, we previously identified a VDAC1 variant which formed dimers/multimers in the membrane, but these were unproductive as the construct failed to scramble lipids ^12^. Thus, the presence of low-activity vesicles in the single-vesicle dataset likely reflects those reconstituted with VDAC1 dimers with suboptimal interfaces. Adding more VDAC1 dimers to a vesicle in the protein/phospholipid ratio regime of our experiments does not necessarily add more active scramblases, further explaining the weak correlation between protein signal and scrambling rate.

To determine the scrambling rate of individual VDAC1 dimers, we analyzed vesicles where Alexa Fluor 647 fluorescence exhibited two-step photobleaching, i.e., vesicles containing a single dimer. These vesicles had Alexa Fluor 647 initial plateau intensities in the range 400–1,000 units, aligning with the average last step height of ∼400 ±150 units (mean ± S.D.) for photobleaching traces (Extended Data Fig. 6) and consistent with our earlier estimate of a threshold of ∼300 units for a monomer. Three exemplary kinetic traces from this group (Fig. 3h; additional examples are shown in Extended Data Fig. 7) illustrate highly active, slightly active, and inactive dimers. The scrambling rate distribution (Fig. 3i) revealed that the top 20% of vesicles, containing optimal dimers, exhibited rates with α >0.003 s⁻¹. Combining this information with vesicle size estimates determined using NBD-PC intensity and dynamic light scattering (Extended Data Fig. 5e) ^31^, and deduced phospholipid content estimates, we obtain a unitary scrambling rate of >1,000 phospholipids per second per VDAC1 dimer for the highly active subpopulation (Fig. 3j).

### Applicability of the single vesicle assay to other scramblases

We tested the general applicability of our assay by investigating another scramblase, the GPCR scramblase opsin ^28,32^. We stoichiometrically labeled bovine opsin using Alexa Fluor 647-maleimide (Extended Data Fig. 8a,b) and reconstituted the labeled protein into NBD-PC-containing LUVs. Ensemble assays using both [S_2_O_4_]^2-^ and BSA revealed the expected scramblase activity (Extended Data Fig. 8c,d)(we note that in these assays, [S_2_O_4_]^2-^ also reports scramblase activity as opsin – unlike VDAC1 – is not a dithionite-permissive channel). The sample was then taken for the single-vesicle scramblase assay. In an initial overview of all NBD fluorescence traces from vesicles associated with one-step Alexa Fluor 647 photobleaching – indicative of the presence of a single opsin molecule in the vesicle – 75% of the vesicles exhibited high scramblase activity, i.e., for these vesicles less than 20% of the initial fluorescence signal remained at the end of the assay (Extended Data Fig. 8e, black traces). The first conclusion that we draw from these data is that opsin monomers have scramblase activity, a point that was previously only inferred from vesicle occupancy statistics ^22,23^. Data fitting using the three-compartment model (Fig. 3g, Extended Data Table 1) revealed that for greater than a third of these vesicles, the scrambling rate constant α was limited by the upper bound of the extraction rate constant γ (Extended Data Fig. 8f). Accordingly, for these vesicles, the calculated unitary transport rate is at least 10,000 lipids per opsin monomer per second (Extended Data Fig. 8g).

## Discussion

We developed a technically accessible, quantitative, high-throughput fluorescence imaging platform to determine the unitary rate of phospholipid scrambling by purified VDAC1 dimers and opsin monomers reconstituted into large unilamellar vesicles. The platform allows the simultaneous measurement of several hundred individual, surface immobilized proteoliposomes, yielding a robust and statistically rich dataset comprising thousands of single vesicles per sample. Our approach overcomes the limitations of traditional ensemble-averaged methods that are typically used to measure lipid scrambling (Extended Data Figs. 3f, 5c, 8c,d).

Although native dimers form due to the high abundance of VDAC1 in the outer mitochondrial membrane ^15^, we purified VDAC1 as a monomer and therefore made use of EGS cross-linking and size-exclusion chromatography to generate and purify a mixture of structurally distinct VDAC1 dimers, each of which is predicted to have a unique protein-protein interface and different scrambling ability. We reconstituted this mixture into liposomes and used stepwise photobleaching (Fig. 3h and Extended Data Fig. 7) to identify vesicles containing a single dimer. By applying a three-compartment mathematical model to analyze lipid exchange kinetics (Fig. 3g) and correlating this with measurements of vesicle diameter and thereby phospholipid content (Extended Data Fig. 5e), we found that individual dimers exhibited a wide range of scrambling rates, from <100 lipids s⁻¹ to >10,000 lipids s⁻¹. The median rate for all active dimers (∼200 lipids s⁻¹) aligns with estimates from ensemble measurements (Fig. 3h-j; ^12^). Poorly active and inactive VDAC1 dimers (Fig. 3h), as well as inactive VDAC1 monomers (Fig. 3b,e, Extended Data Fig. 3f), serve as essential controls, confirming that our reconstitution approach accurately reflects specific protein-mediated scrambling. While bulk assays are often constrained by the need to compare controls that have been treated as similarly as possible to the experimental samples, our approach benefits from an internal control within the sample itself. Specifically, the presence of empty liposomes and inactive dimers within the crosslinked dimer population provides an intrinsic reference.

Our findings support molecular dynamics simulations ^12^, which predict that rapid scrambling occurs at specific VDAC1 dimer interfaces. The distribution of unitary scrambling rates we observed for VDAC1 dimers is somewhat lower than the ensemble-averaged rates reported for Ca²⁺-activated TMEM16 proteins and constitutively active GPCR scramblases ^33^, supported by unitary rates for the latter reported here for opsin (Extended Data Fig. 8g). This suggests that the VDAC1 dimer interface, formed between two β-barrel protomers, may provide an intrinsically slower pathway for lipid transit, yet still sufficient to support mitochondrial membrane biogenesis ^12,34^. With this experimental platform established, future research can focus on (i) characterizing structurally defined VDAC1 dimers generated via site-specific crosslinking to determine their unitary scrambling rates, and (ii) testing different membrane lipid compositions that more closely mimic the mitochondrial outer membrane to evaluate their effect on scrambling activity. We note that in the experiments reported here we used a simple fluid membrane matrix for reconstitution, comprising POPC with 10 mole percentage POPG. This mixture is widely used as it supports the activity of most scramblases. Heterogeneity within a liposome preparation made up of a complex lipid mixture precludes the ready dissection of composition-specific effects in ensemble assays – such analyses are now possible in our single vesicle platform where individual scramblase-containing vesicles can be characterized by properties such as fluidity, or directly in terms of their lipid makeup. Thus, it will now be possible to test precisely the effect of different lipids, including cholesterol, on scramblase activity.

Traditional phospholipid scrambling assays often use [S_2_O_4_]^2-^ to eliminate NBD-fluorescence in the outer leaflet of unilamellar vesicles ^20^, allowing scramblase-mediated lipid translocation to be monitored. However, [S_2_O_4_]^2-^ is unsuitable for measuring VDAC1 dimer scrambling, as it can pass through the VDAC1 pore (Fig. 2d). Nevertheless, within our imaging platform, [S_2_O_4_]^2-^-based assays could be adapted to measure non-pore-forming scramblases such as GPCRs (Extended Data Fig. 8c). In such an approach, vesicles would be reconstituted with long-chain reporter phospholipids labeled at the headgroup with NBD (Fig. 2a) or [S_2_O_4_]^2-^-sensitive ATTO488 ^35^ – these lipids would not desorb during buffer flow but would be probed by introducing [S_2_O_4_]^2-^ into the sample chamber (Fig. 2d), and using the fluorescence decay kinetics to infer scrambling rates. A similar mathematical framework to that in Figure 3g could be applied, with γ representing the pseudo-first-order rate constant for fluorophore reduction by dithionite. Given that γ for [S_2_O_4_]^2-^ reactions is ∼0.07 s⁻¹ (NBD) and ∼0.2 s⁻¹ (ATTO488) ^36^ this method could extend the dynamic range for measuring fast scramblases beyond the γ ∼0.03 s⁻¹ limit of the current setup (Extended Data Fig. 5d, Extended Data Table 1).

Beyond its application to constitutive scramblases such as VDAC1 and opsin, our imaging platform can be directly adapted for studying Ca^2+^-activated scramblases. Thus, the outer leaflet NBD-PC probe can be extracted by buffer flow prior to the start of the experiment, followed by flushing of a Ca^2+^ buffer to activate the scramblase and allow monitoring of scramblase kinetics. Furthermore, our approach can be readily adapted to measure single-protein transport kinetics of ATP-driven flippases, which transport lipids from the exofacial site to the cytosolic site at inherently slow rates due to their reliance on ATP hydrolysis. As these proteins reconstitute preferably ‘inside-out’ with their ATPase domains facing outwards ^18^, their activity could be selectively initiated by introducing ATP into the flow chamber, providing an additional experimental control for functional validation. Here, imaging would begin after extraction of the outer leaflet fluorophores together with a buffer flush supplemented with ATP-Mg or ATP-Na as a control. While bulk measurements require high concentrations of active protein for meaningful readouts, single vesicle microscopy allows selective analysis of the protein-containing subpopulation, enabling more refined and quantitative assessment even in samples with low protein abundance, bypassing the bottleneck of purification. Substrate specificity, dependence on membrane curvature and ultimately the question of whether flippases operate with on and off states similar to proton pumps ^37^ could all be addressed.

In summary, our high-throughput fluorescence imaging platform provides an unprecedented, versatile and scalable tool to measure the unitary rates of phospholipid scramblases, necessary to dissect transport mechanisms and with the potential to study their regulation by membrane factors.

## Methods

### Materials

POPC (#850457), POPG (#840457), 14:0-6:0 NBD-PC (#810122), 16:0-6:0 NBD-PC (#810130), 14:0-6:0 NBD-PE (#810151), 16:0-6:0 NBD-PE (#810153), 16:0-6:0 NBD-PS (#810192), *N*-NBD-DOPE (#810145), C6-NBD-sphingomyelin (#810218) and Biotin-PEG(2000)-DSPE (#880129) were purchased from Avanti Polar Lipids Inc. (Birmingham, AL, USA). Bio-Gel P-6 (#1504130) and Bio-Spin columns (#7326008) were purchased from Bio-Rad (Hercules, CA, USA). Alexa Fluor^TM^ 647 C_2_ Maleimide (#A20347), ethylene glycol bis(succinimidyl succinate) (EGS; #21565), and the Micro BCA™ Protein Assay Kit (#53235) were purchased from Thermo Fisher Scientific (Waltham, MA, USA). PLL(20)-g[3.5]-PEG(2) and PLL(20)-g[3.5]-PEG(2)/PEG(3.4)-biotin(20%) were obtained from SuSoS (Dübendorf, Switzerland). All other chemicals and reagents, including fatty acid-free bovine serum albumin (#126609), Tetramethylrhodamine-5-maleimide (#94506), sodium dithionite (#157953), and the zwitterionic detergent N,N-Dimethyl-n-dodecylamine N-oxide (LDAO; #40236) were obtained from Sigma-Aldrich, unless otherwise stated.

### VDAC1 purification and fluorescence labeling

PCR-based site-directed mutagenesis was used to generate a human VDAC1 construct (C127S, A170C, C232S) with a single cysteine residue facing the protein pore. This construct is referred to as VDAC1-A170C or simply VDAC1. VDAC1 was expressed in BL21 (DE3)-omp9 *E. coli* and purified from inclusion bodies as described previously, using Ni-NTA affinity chromatography followed by size exclusion ^12,38^. For fluorescence labeling, 340 µg of purified VDAC1 (100 µL of a 100 µM VDAC1 solution in 20 mM Tris-HCl, pH 8.0, 150 mM NaCl, 0.1 % (w/v) LDAO) was reacted with a 10-fold molar excess of Alexa Fluor 647 C_2_ Maleimide (from a 10 mM stock solution in DMSO) or mock-treated with DMSO. The sample was incubated (2 h, 40°C, 650 rpm) in an Eppendorf Thermomixer, after which the labeling reaction was stopped by incubating for a further 15 min with dithiothreitol, added from a 1 M stock solution to give a 20-fold molar excess over Alexa Fluor 647 C_2_ Maleimide. The sample was then subjected to three rounds of buffer exchange using Bio-Gel P-6 resin (1 mL bed volume in Bio-Spin columns equilibrated in crosslinking buffer (10 mM MOPS-KOH pH 7.0, 100 mM KCl, 0.05% LDAO)). Protein concentration after buffer exchange was determined via absorbance at 280 nm (ε_280_ = 32,500 M^-1^ cm^-1^) as well as with the Micro BCA™ Protein Assay Kit, with a typical recovery of ∼40%. To quantify labeling efficiency, equivalent amounts of Alexa Fluor 647 C_2_ Maleimide-treated and unlabeled VDAC1 were reacted with tetramethylrhodamine-5-maleimide (added from a 0.2 mM stock in DMSO in 10-fold molar excess over VDAC1) following the same protocol. As tetramethylrhodamine is smaller than Alexa Fluor 647 it is expected to react quantitatively with solvent-accessible cysteines. Rhodamine-labeled samples were analyzed by SDS-PAGE followed by visualization and labeling quantification using the Bio-Rad ChemiDoc™ MP Imaging System. This protocol indicated that that Alexa Fluor 647 labeling was 97 ± 3% efficient (mean ± SD, n = 2) as ∼3% of the sample could be additionally labeled with rhodamine.

### VDAC1 crosslinking

Alexa Fluor™ 647-labeled VDAC1 was treated with the homobifunctional amino-reactive crosslinker EGS as described ^12^ to generate a mixture of crosslinked species (CL). A mock-treated sample was prepared in parallel (NCL, non-crosslinked). Dimers and multimers in the CL sample were resolved from monomers by size exclusion chromatography using a Superdex 200 Increase 10/300 GL column run in 20 mM Tris-HCl, pH 8.0, 150 mM NaCl, 0.1 % (w/v) LDAO. Fractions were analyzed by SDS-PAGE, and in-gel Alexa 647 fluorescence was used to identify the most enriched dimer fraction (pCL, purified crosslinked).

### Preparation of large unilamellar vesicles (LUVs)

LUVs composed of POPC, POPG (9:1, molar ratio), 0.5 mol% 14:0-6:0 NBD-PC and 0.1 mol% Biotin-PEG(2000)-DSPE were prepared in reconstitution buffer (10 mM MOPS/KOH pH 7.0, 100 mM KCl) following a previously described protocol ^12^. For some experiments 0.5 mol% *N*-NBD-DOPE or other NBD-lipids as indicated were used instead of 14:0-6:0 NBD-PC. As NBD-PC is present in both leaflets of the membrane, the LUVs are considered symmetrically labeled. The LUVs have a typical phospholipid concentration of 3.5 mM, and a hydrodynamic diameter of 169 ± 4 nm and polydispersity of 8.7 ± 5.3% (intensity-weighted size distribution, mean ± SD, n=6), as determined by Dynamic Light Scattering using an Anton Paar Litesizer™ 500 Particle Analyzer. To prepare vesicles with 14:0-6:0 NBD-PC located exclusively in the outer leaflet (asymmetrically labeled vesicles), NBD-PC was left out of the original liposome preparation and added afterwards. Briefly, 12 µg of 14:0-6:0 NBD-PC was dissolved in 1 µL ethanol and 9 µL reconstitution buffer; 2 µL of this solution was added to 50 µL unlabeled liposomes, and the sample was diluted with reconstitution buffer to a total volume of 100 µL. The sample was incubated on ice for 10 min to ensure complete outside labeling and subsequently used for measurements.

### VDAC1 reconstitution into LUVs, and ensemble assays of scrambling

Non-crosslinked (NCL), cross-linked (CL) and purified crosslinked (pCL) Alexa Fluor™ 647-labeled VDAC1 dimers were reconstituted into NBD-PC-containing LUVs using a detergent-destabilization protocol as described ^12,38^. For reconstitution, 800 µL LUVs and 10 µg VDAC1 preparations were used, unless otherwise stated. Protein-free liposomes were prepared in parallel. The resulting vesicles had a phospholipid concentration of 2.38 ± 0.05 mM (n=4), a hydrodynamic diameter of 208 ± 10 nm and polydispersity of 13 ± 6% (mean ± SD, n=12). The specific values for the pCL-reconstituted vesicles were 203 ± 9 nm (diameter) and 13.4 ± 9.3% (polydispersity). The vesicles were assayed for channel activity using sodium dithionite (Na_2_S_2_O_4_), and scramblase activity using fatty acid-free bovine serum albumin (BSA) using fluorescence-based ensemble assays as previously described ^12^. Briefly, 50 µL of vesicles were diluted to 2.5 mL in buffer (50 mM HEPES, 150 mM NaCl, pH 7.4) in a cuvette equipped with a magnetic stir bar. NBD fluorescence (λ_ex_=470 nm, λ_em_=530 nm) was monitored under continuous stirring using a fluorescence spectrometer. After signal stabilization, dithionite (final 20 mM; from a freshly prepared 1 M stock in 0.5 M Tris base, pH 10) or fatty acid-free BSA (final 15 mg/mL, from a 75 mg/mL stock in 50 mM HEPES, 150 mM NaCl, pH 7.4) were added, and fluorescence was monitored every second for up to 10 min. Dithionite eliminates NBD fluorescence by chemically reducing the fluorophore, whereas BSA binds to desorbed NBD-PC resulting in fluorescence quenching. Time-dependent fluorescence loss was analyzed after normalization to the initial fluorescence signal.

### Opsin purification, fluorescence labeling, reconstitution into LUVs and ensemble assays of scrambling

Opsin was purified from bovine retina as described previously ^17^ and labeled with Alexa Fluor 647 C_2_ maleimide by adding 2.5 µL of 1 mM dye (in DMSO) to 2.5 µM opsin in 100 µL labeling buffer (50 mM Hepes, pH 7.5, 150 mM NaCl, 0.1% n-dodecyl-β-maltoside). The sample was incubated overnight at 4°C, followed by three rounds of buffer exchange using Biogel P6 resin (1 ml bed volume, pre-equilibrated in labeling buffer). Spectrophotometric analysis confirmed a 1:1 labeling stoichiometry (extinction coefficients: 270,000 M⁻¹ cm⁻¹ for Alexa Fluor 647; 81,200 M⁻¹ cm⁻¹ for opsin), consistent with previous reports ^39,40^. Under our labeling conditions the C316 residue is likely to be the principal site of fluorescence modification ^40^. SDS-PAGE with fluorescence imaging and Coomassie staining revealed a single ∼35 kDa band. Approximately 0.4% of the fluorescence of the protein band was found as free dye running at the dye-front of the gel, indicating that the desalting protocol to remove free dye was efficient. The fluorescently labeled opsin (2.5 and 5 µg aliquots) was reconstituted into preformed vesicles (400 µl of a 3.6 mM stock) as described above for VDAC. Ensemble scramblase assays were performed as described above using dithionite and BSA, both of which report scrambling for opsin (this is in contrast to VDAC, where only BSA reports scrambling). Reconstituted vesicles had a phospholipid concentration of 2.8 mM and a hydrodynamic diameter of 176.6 ± 5.3 nm with 13.2 ± 6.9% polydispersity (mean ± SD, n=6).

### Assembly of the sample chamber for single vesicle imaging

The process is illustrated in Extended Data Figure 1. Glass coverslips (26 x 76 mm, thickness #1.5, Thermofisher) were sonicated twice for 15 min in 2 % (v/v) Hellmanex III solution, thoroughly rinsed with ddH_2_O, then sonicated in methanol and stored in fresh methanol. After drying with compressed air, the coverslips were ozone-cleaned for 15 min using a Novascan PSD Pro UV Ozone system (Novascan technologies, Boone, IA, USA). A bottomless 6-channel sticky slide (Sticky-Slide VI 0.4, ibidi GmbH, Gräfelfing, Germany) was adhered to the cleaned coverslip (channels: 17 x 3.8 x 0.4 mm, 30 µL channel capacity, 60 µL per well). The channels were flushed with 60 µL of 1 mg/mL Biotinyl-PLL-PEG / PLL-PEG (1:1,000) in PBS (150 mM NaCl, 8 mM Na_2_HPO_4_, 2 mM KH_2_PO_4_, pH 7.0). After 20 min, the channels were flushed 5 times with 60 µL of reconstitution buffer to remove unbound material. Biotin groups were saturated by introducing 60 µL of 0.025 g/L NeutrAvidin in reconstitution buffer, incubating for 20 min, then flushing 5 times with 60 µL of reconstitution buffer to remove unbound NeutrAvidin. The assembly was used immediately for microscopy or stored at 4°C for up to a week.

### TIRF microscopy

Single vesicle imaging was done using an inverted TIRF microscope (IX83, Olympus, Hamburg, Germany) with a 100x oil-immersion objective (1.5 NA; Olympus), an Orca-flash 4.0 sCMOS camera (Hamamatsu Photonics K.K., Hamamatsu, Japan), and a quad band emission filter cube (DAPI/FITC/CY3/CY5). Lasers were aligned and refocused before each session. NBD fluorophores were excited using a 488 nm laser (green channel) at 10% intensity (18.3 mW/cm^2^) with 25 ms exposure time and the Alexa 647 protein marker was excited with a 640 nm laser (red channel) at 15% intensity (20.2 mW/cm^2^) and 200 ms exposure time. Both laser angles were adjusted to 200 nm penetration depth. To avoid uncertainty at the edges of the illuminated area, the central 1,024 x 1,024-pixel region, corresponding to 66.56 × 66.56 μm in 16-bit range, was used for analysis, resulting in a pixel size of 65 × 65 nm. The IX-ZDC z-drift compensator (Olympus) maintained the focal plane during long-term imaging.

### Single vesicle phospholipid scramblase assay

Measurements were made at ambient temperature unless otherwise stated. Tubing (Masterflex Tygon E-lab L/S 13; Bohlender PTFE 0.5 x 1.0 mm, I.D x O.D) was connected to the wells of the channel slide for buffer flow using a peristaltic LabV1 pump with an AMC2 head (Drifton, Hvidovre, Denmark)(Extended Data Figure 1). The tubing and channel were flushed with reconstitution buffer for 1 min (500 µL/min), followed by flushing the buffer-diluted sample (1:1,500) for 1 min at 500 µL/min. Buffer flow was paused for 1 min to enable vesicle immobilization. A single image was then captured before resuming buffer flow (500 µL/min) concomitantly with time-lapse imaging. Using the CellSens experiment manager (Version 3.2; Olympus), one image was taken every 90 s in the lipid channel, with up to a total of 10 images in 15 min. Subsequently, the same frame was repeatedly imaged 1,250 times in the red channel for photobleaching step analysis.

### Single vesicle dithionite assay to determine vesicle occupancy

The experimental setup was similar to the single vesicle scramblase assay. After sample immobilization, a 5-min buffer flush (500 µL/min) removed all accessible outer leaflet short-chain NBD lipids. Using a multi-position imaging tool, 12 frames were analyzed simultaneously as described previously ^18^ and three imaging rounds, each capturing three images per frame in both channels, were performed. After the first round, reconstitution buffer containing 10 mM dithionite (from a 1 M stock in 0.5 M Tris pH 10) was flushed through the channel for 1 min followed by 2 min incubation. After the second round, alamethicin (10 μM, prepared in 10 mM dithionite) was flushed for 1 min with an additional 2 min incubation to ensure completion of alamethicin-induced permeabilization and dithionite-mediated elimination of NBD fluorescence before the final imaging round.

### Data analysis

Recorded .vsi/.ets files were converted to TIFF format in FIJI (2.9.0/1.52v) using bio-formats reader with batch processing without interpolation. TIFF files were analyzed with a custom Python algorithm based on a previously described anaylsis ^18^. Particle centers and corresponding intensity maxima were tracked in the lipid channel TIFF stack using *trackpy* ^41^, with local background-corrected intensities extracted using *photutils* and *astropy* ^42^. Particle coordinates determined from the lipid channel were used to extract protein channel signals from the same regions. Lipid and protein thresholds were optimized using a protein-free sample. As described in detail in the Results section, lipid fluorescence of scramblase-containing proteoliposomes decays rapidly at first, corresponding to the loss of outer leaflet lipids to the flow of buffer, and then more slowly as lipids from the inner leaflet are scrambled out and lost to buffer flow in turn. Vesicles with no decrease in lipid signal upon buffer flow or those where the signal decreased sharply (indicative of detachment from the chamber) were excluded from the analysis. The extraction rate constant (γ), i.e., the rate of loss of outer leaflet lipids to buffer flow, was determined as 0.0217 ± 0.01 s^-1^ by fitting traces from protein-free liposomes (Fig. 3a) to a mono-exponential function. When fitting fluorescence decay traces for scramblase active-proteoliposomes based on a three compartment model adapted from ^27^ using *scipy* (see Supplementary information), three models with 2-3 variables were tested. Based on the Akaike and Bayesian Information Criteria (AIC, BIC; Extended Data Table 1), Model 1 was selected for further analyses where γ was constrained to the bounds 0.0117-0.0317 s^-1^ and the initial ratio of outer to inner leaflet NBD-lipid was constrained to the bounds 0.40-0.60. To estimate vesicle size based on lipid signal, the lipid signal intensity distribution average was normalized by the sample average determined by DLS (Extended Data Fig. 5d) as described previously ^18,31^.

## Data availability

Source data are provided with this paper.

## Code availability

Home written analysis code is described in detail in the Methods. Code is available upon reasonable request.

## Acknowledgements

Figures were created with BioRender software (https://biorender.com). We thank Joshua Levitz for comments. This work was supported by the National Institutes of Health (grant no. R01GM146011 to A.K.M.), Canadian Institutes of Health Research Operating Grants PJT-159464 and PJT-195648 (to O.P.E.). and Deutsche Forschungsgemeinschaft (grant no. VE 1674/1-1 to S.V.; grant nos. GU 1133/13-1 and INST 213/985-1 to T.G.P.).

## Author contributions

S.V., A.K.M. and T.G.P. conceived the study. K.M.M. contributed the mathematical solution of the three-compartment model and the associated fitting software. G.I.D. and F.N. prepared and characterized VDAC1 proteoliposomes. T.M. and O.P.E. provided purified bovine opsin. I.M. measured the critical bilayer concentration of NBD-PC and prepared and characterized opsin proteoliposomes. S.V. wrote software for single-vesicle measurements and performed and analyzed single vesicle experiments the experiments under the supervision of A.K.M. and T.G.P.. S.V. prepared the initial draft of the manuscript, including the figures. The final manuscript was prepared by S.V., A.K.M. and T.G.P. and discussed with all authors.

## Additional information

**Extended data** are provided with the manuscript.

**Supplementary information** is provided with the manuscript.

---

**Extended Data Figure 1.**
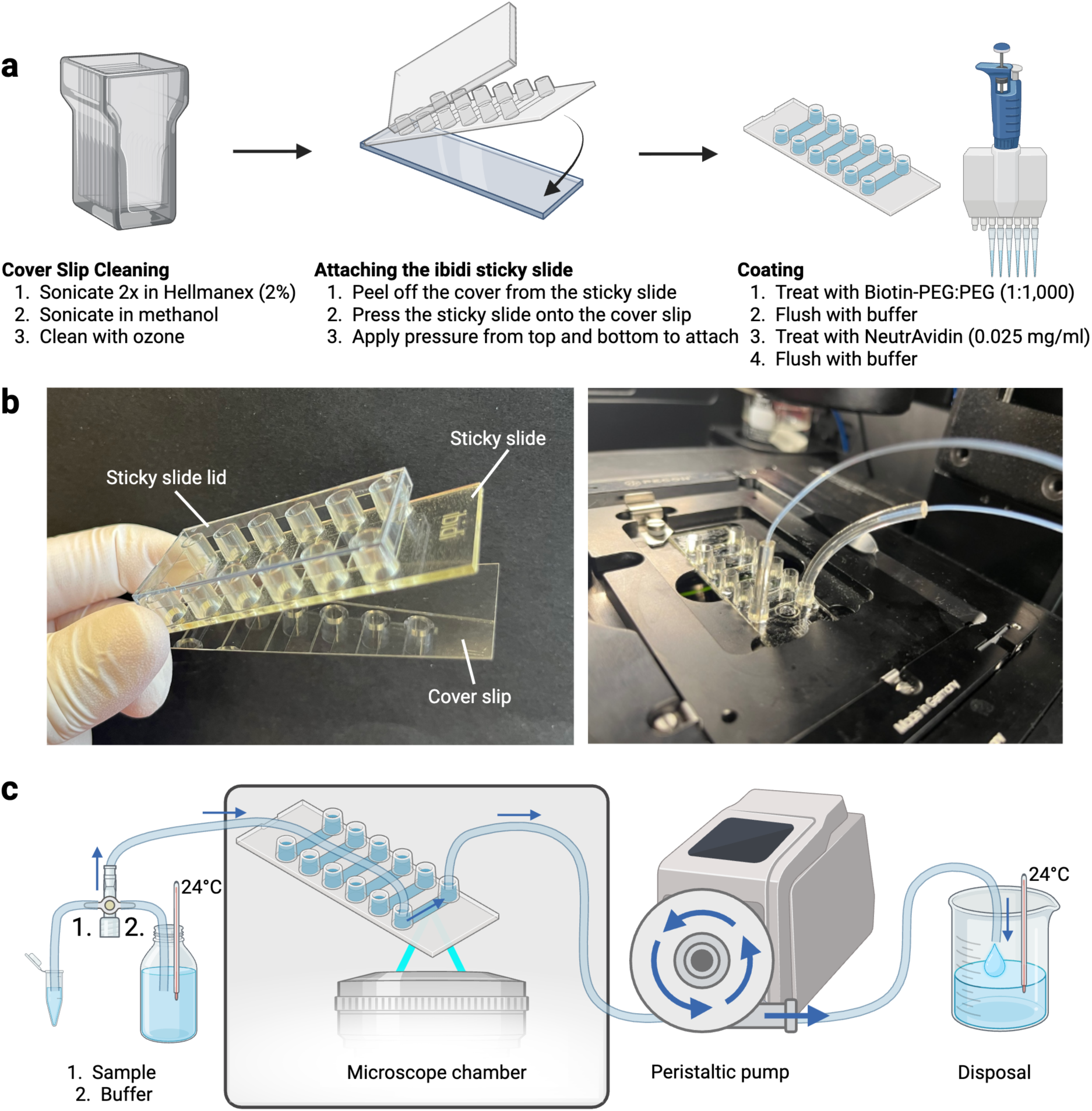
Assembly of the sample chamber for single vesicle imaging. **a**, Glass cover slips (26 x 76 mm) are sequentially cleaned by sonication in 2% Hellmanex solution and washed in ddH₂O, followed by sonication in MeOH. After drying using compressed air, the cover slips are ozone cleaned. Next, a 6-channel ibidi sticky slide is pressed onto each cover slip to form six independent channels. Channel surfaces are passivated with a 1:1,000 mixture of biotin-PEG and PEG, introduced via the channel ports. After flushing five times with buffer using a multipipette (the spacing of the pipette tips matches that of the ibidi chamber), NeutrAvidin is applied followed by an additional five buffer flushes, after which the sample chamber is ready for downstream use. **b**, Photograph of the sticky slide assembly (left) and the sample chamber mounted on the microscope stage (right). **c**, Tubing is connected to the channel in the foreground (as also seen in the right image in panel **b**). A peristaltic pump (500 µl/min) is used to introduce the sample into the channel, followed by buffer flushing to start the assay. Channel temperature is estimated by monitoring the temperature of the buffer in the reservoir and waste container.

**Extended Data Figure 2.**
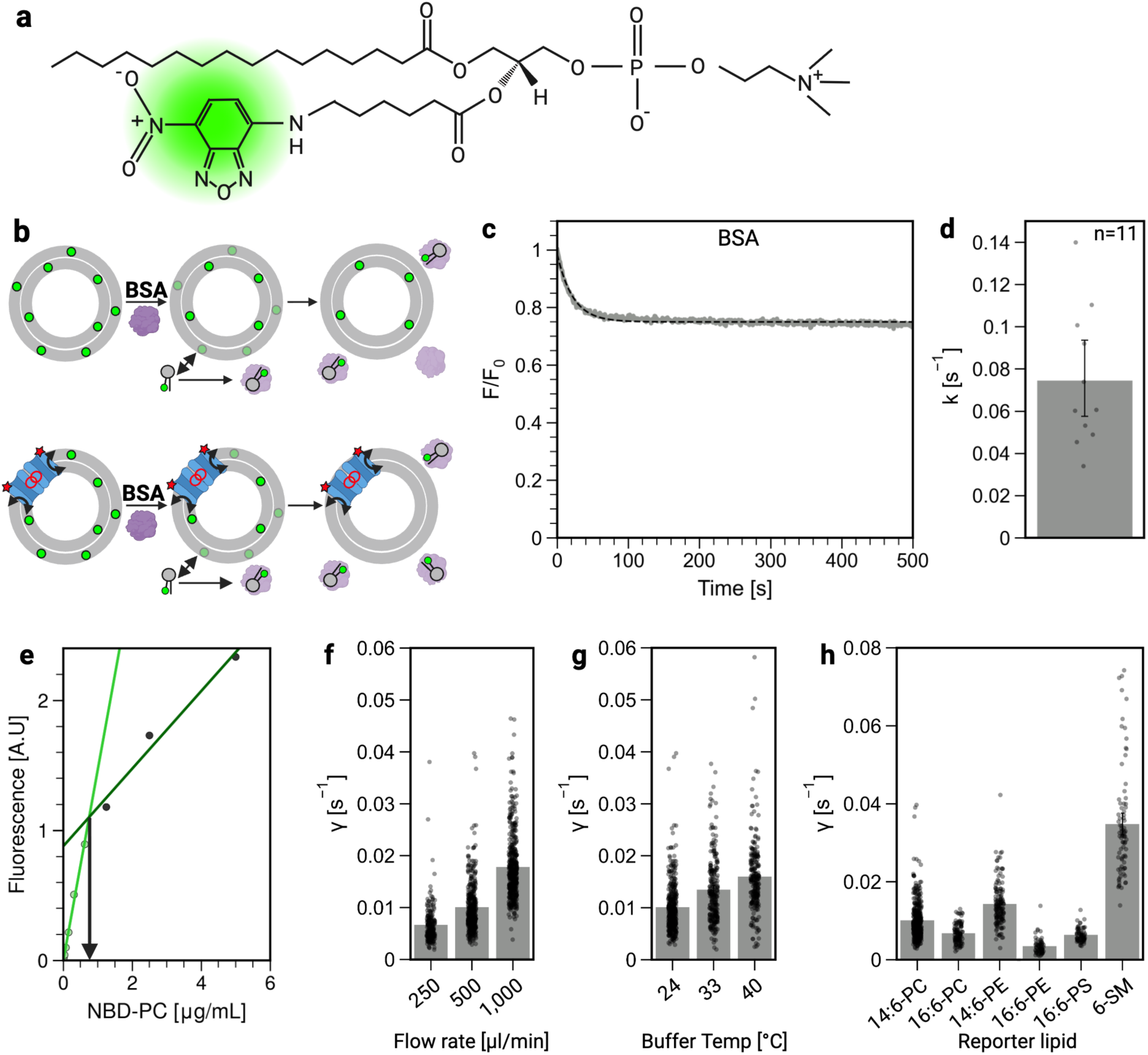
Determination of lipid extraction rate. **a**, Structure of 14:0-6:0 NBD-PC, with the NBD fluorophore highlighted in green. **b**, Scheme of ensemble assay using BSA. Fatty acid-free BSA is added in excess to a suspension of vesicles in a stirred cuvette. NBD-PC molecules in the outer leaflet of the vesicles partition into the aqueous phase, where they are captured by BSA. As the fluorescence quantum yield of NBD is ∼2-fold lower when NBD-PC is bound to BSA versus located in the membrane, a drop in fluorescence is observed. In scramblase-equipped vesicles, NBD-PC in the inner-leaflet is translocated to the outer leaflet and subsequently captured by BSA. This results in a greater drop in fluorescence for scramblase-equipped vesicles compared with protein-free liposomes. This assay enables real-time measurement of lipid translocation and extraction kinetics in a bulk population of vesicles. **c**, Time-course analysis of the BSA assay performed on 14:0-6:0 NBD-PC-containing protein-free liposomes. The data are expressed as normalized fluorescence intensity relative to the initial value, following the addition of BSA at time=0. The desorption process is monitored by measuring the decrease in fluorescence over time, which reflects the transfer of NBD-PC from the liposomal membrane to BSA. **d**, Kinetic analysis of NBD-PC desorption from protein-free liposomes. Fluorescence decay traces were fitted to a mono-exponential decay function (for example, dashed line in panel **c**), yielding a desorption rate constant of *k* = 0.07 s⁻¹ for NBD-PC (*n* = 11). Error bar represents the 95% confidence interval. **e**, Fluorescence intensity as a function of NBD-PC concentration in an aqueous environment, used to determine its solubility threshold (0.75 µM) as the point at which monomeric NBD-PC self-associates, resulting in a change in fluorescence quantum efficiency. This point was determined as the intersection of two lines corresponding to linear fits of the two portions of the graph. **f-h**, LUVs were prepared with 14:0-6:0 NBD-PC (panels **f** and **g**) or with various fluorescent NBD lipids as indicated (panel **h**). The vesicles were immobilized and an initial TIRF microscopy image was acquired, before initiating buffer flow resulting in decay of the fluorescence signal due to desorption of fluorescent lipids from the outer leaflet and their elimination from the chamber. For each vesicle, fluorescence was normalized to the initial intensity, and the time-dependent loss of fluorescence was analyzed via a mono-exponential function to determine the desorption rate constant γ. Different flow rates (**f**) and buffer temperatures (**g**) were tested with 14:0-6:0 NBD-PC-containing LUVs. In panel **h**, LUVs containing various NBD-lipids were analyzed under standard settings (500 µl/min, 24°C). **f-g**: Statistical significance was assessed using one-way ANOVA with post-hoc Tukey’s test with all pairwise comparisons being significant with p < 0.001 in **f** and **g**. In **h**, statistical significance was p < 0.001 for all pairwise comparisons except for 16:6 PC-16:6 PE which is p < 0.01, and 16:6 PC-PS and 16:6 PE-PS which are not significant. For panels **f–h**, the filter settings used to discard unusable traces (e.g., corresponding to vesicles that detached during imaging, or those where there was no change in fluorescence) were adjusted to include lower extraction rates, and identical settings were applied across all samples to enable direct comparison.

**Extended Data Figure 3.**
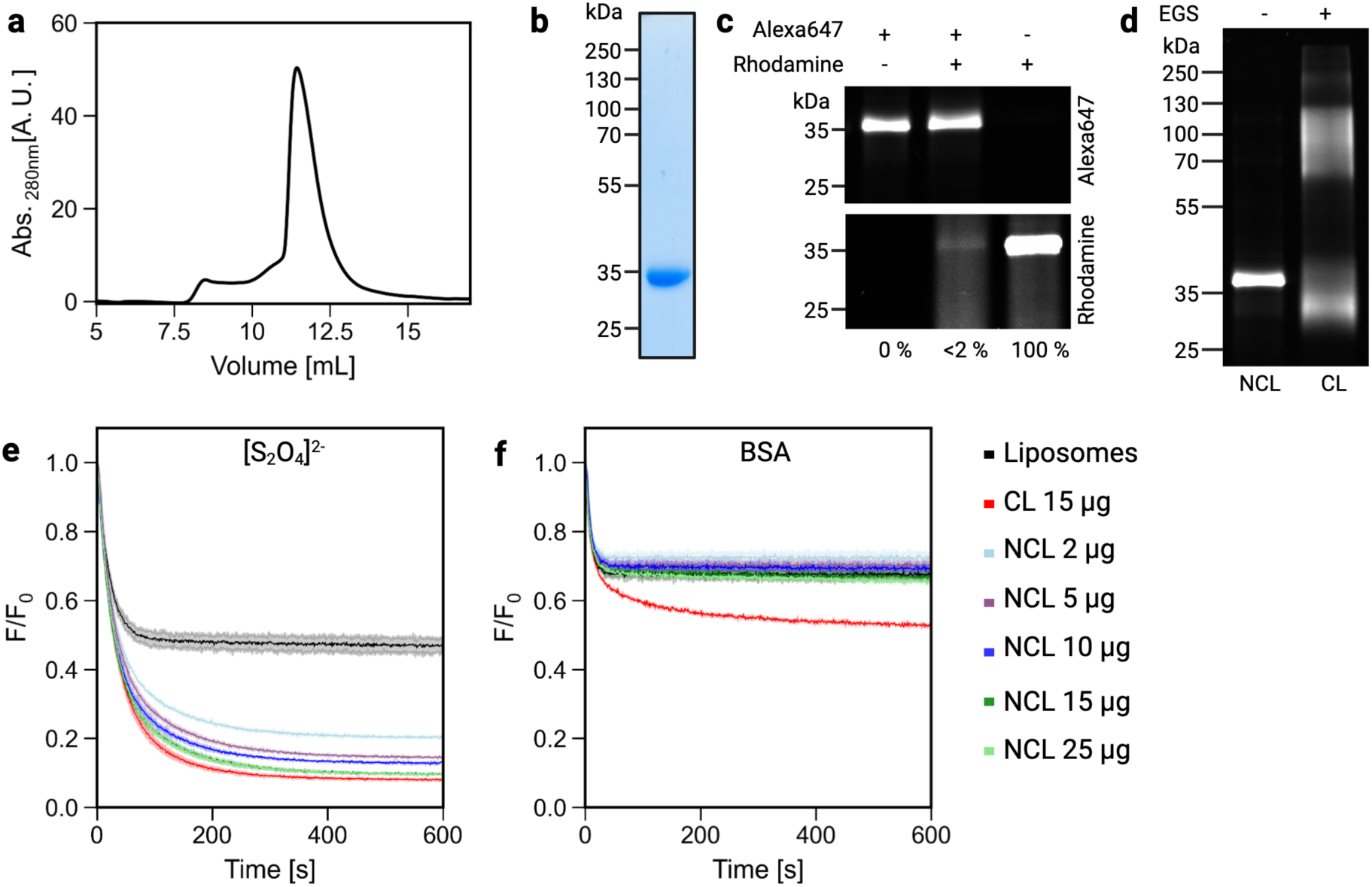
VDAC1-A170C purification, fluorescence labeling and crosslinking. **a**, Size exclusion chromatography profile of purified VDAC1-A170C. **b**, Coomassie-stained SDS-PAGE of peak fraction. **c**, Purified VDAC1-A170C was labeled via Alexa Fluor 647-maleimide (lane 1) and subsequently labeled with rhodamine-maleimide to detect unreacted cysteines (lane 2). In comparison with a sample that was directly labeled with r<u>h</u>odamine (lane 3), only 1.3 % of the Alexa Fluor 647-labeled preparation was susceptible to rhodamine labeling in the example shown, indicating high efficiency (>97 ± 3%; mean ± SD, n = 2) of the initial Alexa Fluor 647 conjugation. An SDS-PAGE gel is shown, imaged to visualize Alexa Fluor 647 (top panel) and rhodamine (lower panel). **d**, Alexa Fluor 647-labeled VDAC1 was either mock-treated with DMSO (non-crosslinked, NCL) or crosslinked (CL) using EGS to generate dimers/multimers. An SDS-PAGE gel is shown, imaged to visualize Alexa Fluor 647. Monomeric VDAC1 in the EGS-treated sample migrates faster than in the control sample because of intramolecular crosslinking. **e, f**, 14:0-6:0 NBD-PC-containing protein-free liposomes or proteoliposomes reconstituted with the indicated amounts of either non-crosslinked (NCL) or crosslinked (CL) Alexa Fluor 647-labeled VDAC1 were subjected to dithionite ([S_2_O_4_]^2-^) reduction or BSA back extraction assays in a cuvette (the dithionite assay detects vesicles reconstituted with a VDAC1 channel, whereas the BSA assay detects vesicles with scramblase activity). **e**, Time-dependent decay of 14:0-6:0 NBD-PC fluorescence (normalized to the starting value) in protein-free liposomes and VDAC1 containing proteoliposomes upon dithionite addition at t=0 s; **f**, Time-dependent decay of 14:0-6:0 NBD-PC fluorescence (normalized to the starting value) in protein-free liposomes and VDAC1 containing proteoliposomes upon BSA addition at t=0 s.

**Extended Data Figure 4.**
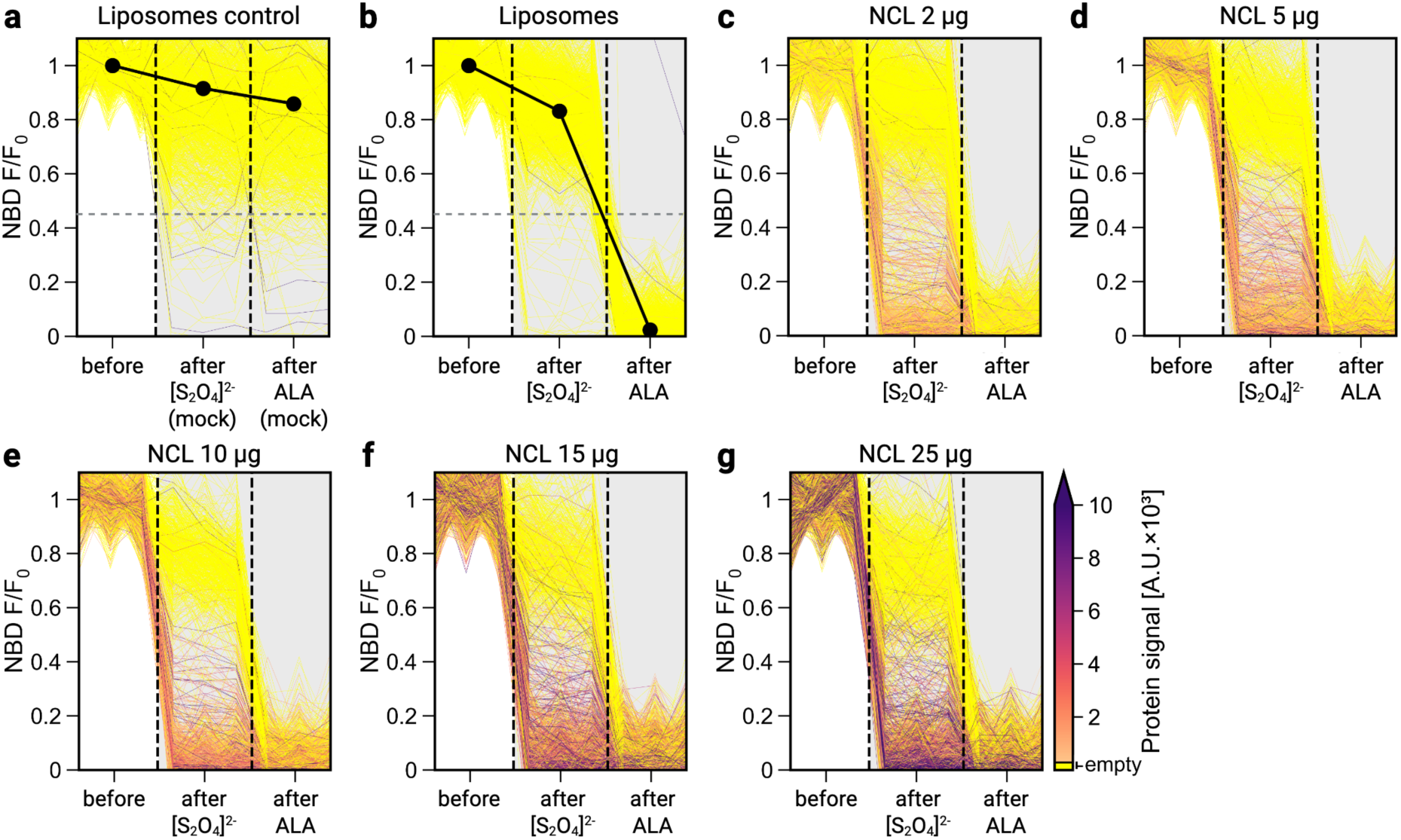
Single vesicle analysis of VDAC1 occupancy. 14:0-6:0 NBD-PC-containing protein-free liposomes or proteoliposomes with indicated amounts of non-crosslinked (NCL) Alexa Fluor 647-VDAC1 were immobilized as described in Figure 1. NBD-PC from the outer leaflet of the vesicles was removed flushing the chamber with buffer for 5 min. Then three ‘before’ images were taken in the lipid channel, with the average set as F_0_. Next, the chamber was flushed with 10 mM sodium dithionite ([S_2_O_4_]^2-^) for 1 min followed by 2 min incubation at the first dashed line in each plot. At this point ‘after [S_2_O_4_]^2-^’ images were taken. Subsequently, alamethicin (ALA) in 10 mM [S_2_O_4_]^2-^ was flushed for 1 min with 2 min incubation (second dashed line), and then ‘after ALA’ images were acquired to assess further loss of NBD-PC fluorescence due to vesicle permeabilization and dithionite access to the inner leaflet pool. Each trace represents the change in NBD-PC fluorescence intensity from single vesicles, with protein signals color-coded according to the colormap shown to the right of panel g. Protein-free liposomes were measured either without additions to estimate photobleaching and general signal stability (**a**) or with additions (**b**). **a**,**b**, Horizontal grey dashed lines indicate the set threshold of 0.45 used to distinguish vesicles where the inner leaflet pool of NBD-PC is exposed to dithionite, either on account of a VDAC channel or by the addition of ALA. The average value of F/F_0_ is shown for each of the panels (black circle). Fluorescence traces for single proteoliposomes containing non-crosslinked (NCL) VDAC1 are shown in (**c-g**).

**Extended Data Figure 5.**
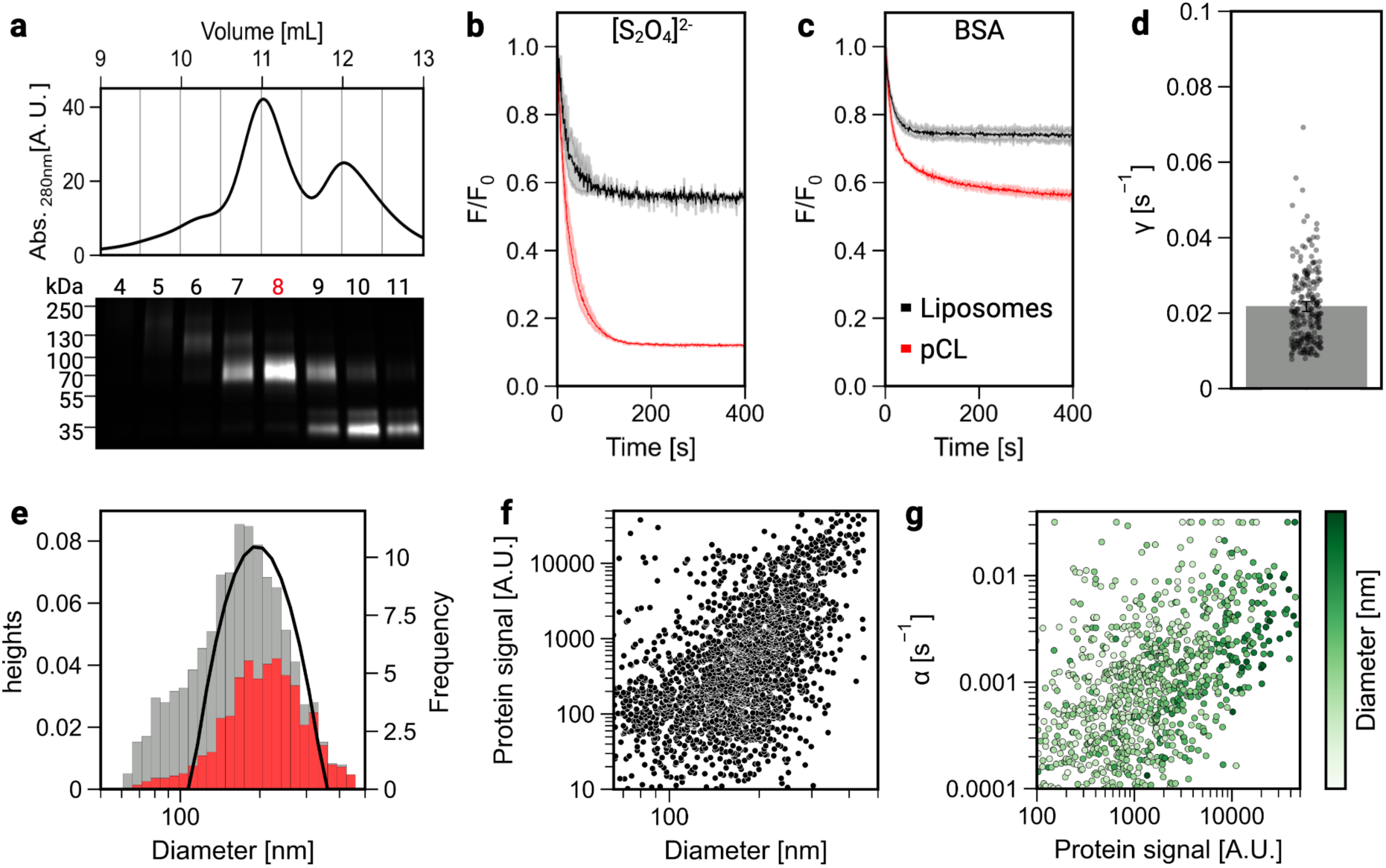
Characterization of proteoliposomes reconstituted with purified VDAC1 dimers. **a**, Size-exclusion chromatography profile of EGS-crosslinked Alexa Fluor 647-labeled VDAC1 (top panel) and an Alexa Fluor 647 fluorescence image of an SDS-PAGE gel of collected fractions (bottom panel). Fraction 8, highly enriched in purified cross-linked (pCL) VDAC1 dimers was used for reconstitution. **b, c**, Cuvette-based assays of NBD-PC-containing protein-free liposomes and pCL proteoliposomes using sodium dithionite ([S_2_O_4_]^2^) (**b**) and BSA (**c**) added at t=0 s. **d**, Fluorescence decay traces from single vesicle measurements of protein-free liposomes reported in Fig. 3a were fitted to a mono-exponential decay function. The desorption rate constant (γ) was determined for each individual vesicle (dots), yielding an average desorption rate of 0.0217 ± 0.010 s⁻¹ (mean ± S.D., n=254). Error bar represents the 95% confidence interval. **e**, Dynamic light scattering intensity-weighted size distribution (solid line) indicating an average vesicle diameter of 203 nm. The lipid fluorescence intensity distribution histogram of the sample obtained via TIRF microscopy, normalized to the DLS data, with vesicles categorized as protein-containing (red) or empty (grey). **f**, Correlation analysis of protein signal intensity (Alexa Fluor 647) versus vesicle diameter (deduced from lipid signal intensity (NBD)). **g**, Values of the scrambling rate constant (alpha, see Fig. 3g), derived from single-vesicle traces fitted to the three-compartment kinetic model using Model 1, plotted against initial protein fluorescence signal of the vesicles. Data points are color-coded by vesicle diameter, based on lipid signal normalized via DLS.

**Extended Data Figure 6.**
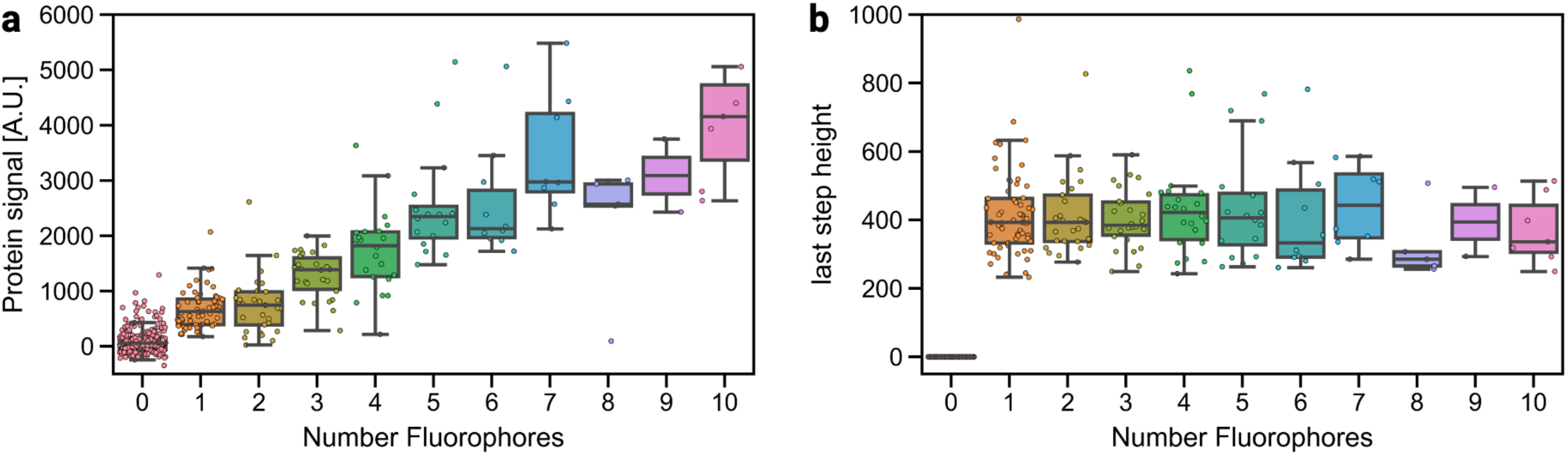
Intensity of single Alexa Fluor 647 fluorophores determined by bleaching step analysis. Proteoliposomes containing Alexa Fluor 647-labeled pCL VDAC1 were immobilized and subjected to photobleaching step analysis using TIRF microscopy. **a**, Initial protein fluorescence signal of each vesicle is plotted against the total number of detected fluorophores per vesicle, as determined by discrete photobleaching steps. As the fitting success is reduced with increasing fluorophore number, the graph is limited to vesicles with 10 fluorophores. **b**, Intensity of the final photobleaching step, categorized by fluorophore count per vesicle. Single-fluorophore emission ranges from ∼250–550 A.U.

**Extended Data Figure 7.**
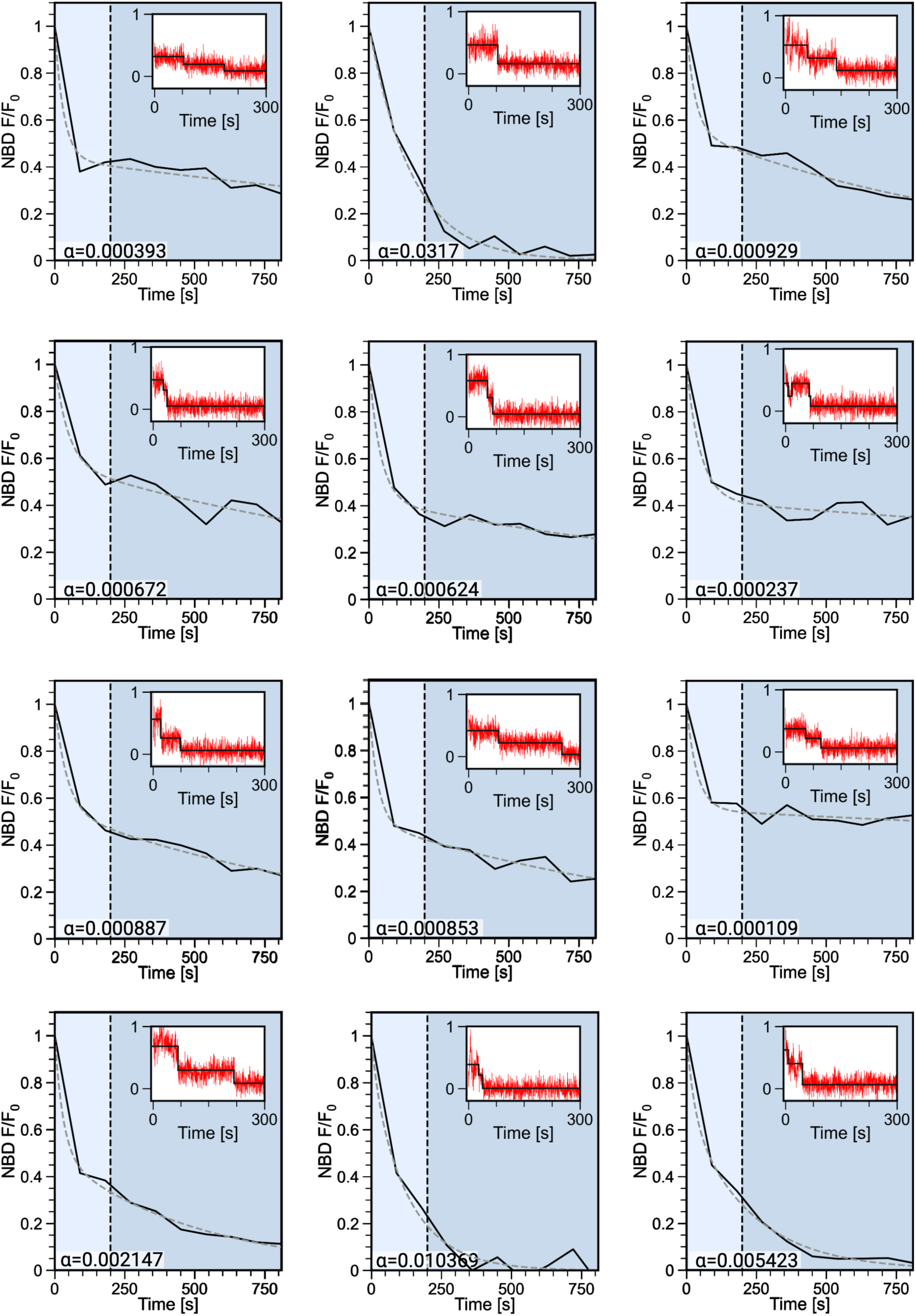
Exemplary fluorescence decay traces for proteoliposomes containing one pCL VDAC1 dimer. 14:0-6:0 NBD-PC proteoliposomes containing pCL Alexa Fluor 647-labeled VDAC1 were immobilized and subjected to TIRF microscopy as described in Fig. 1. Imaging was started just before buffer flow (t=0 min). For each individual liposome, NBD fluorescence intensity is shown as a normalized line plot relative to the initial fluorescence. The light blue section of each graph demarcates the initial 200 s period of buffer flow during which all outer leaflet NBD-PC is expected to be eliminated (Fig. 2c). The scrambling rate constant α was determined using the Model 1 version (Extended Data Table 1) of a three-compartment model (fit shown as dashed line) and is displayed in the lower-left corner (units s^-1^). The *Inset* shows the corresponding photobleaching step analysis for the protein fluorophores.

**Extended Data Figure 8.**
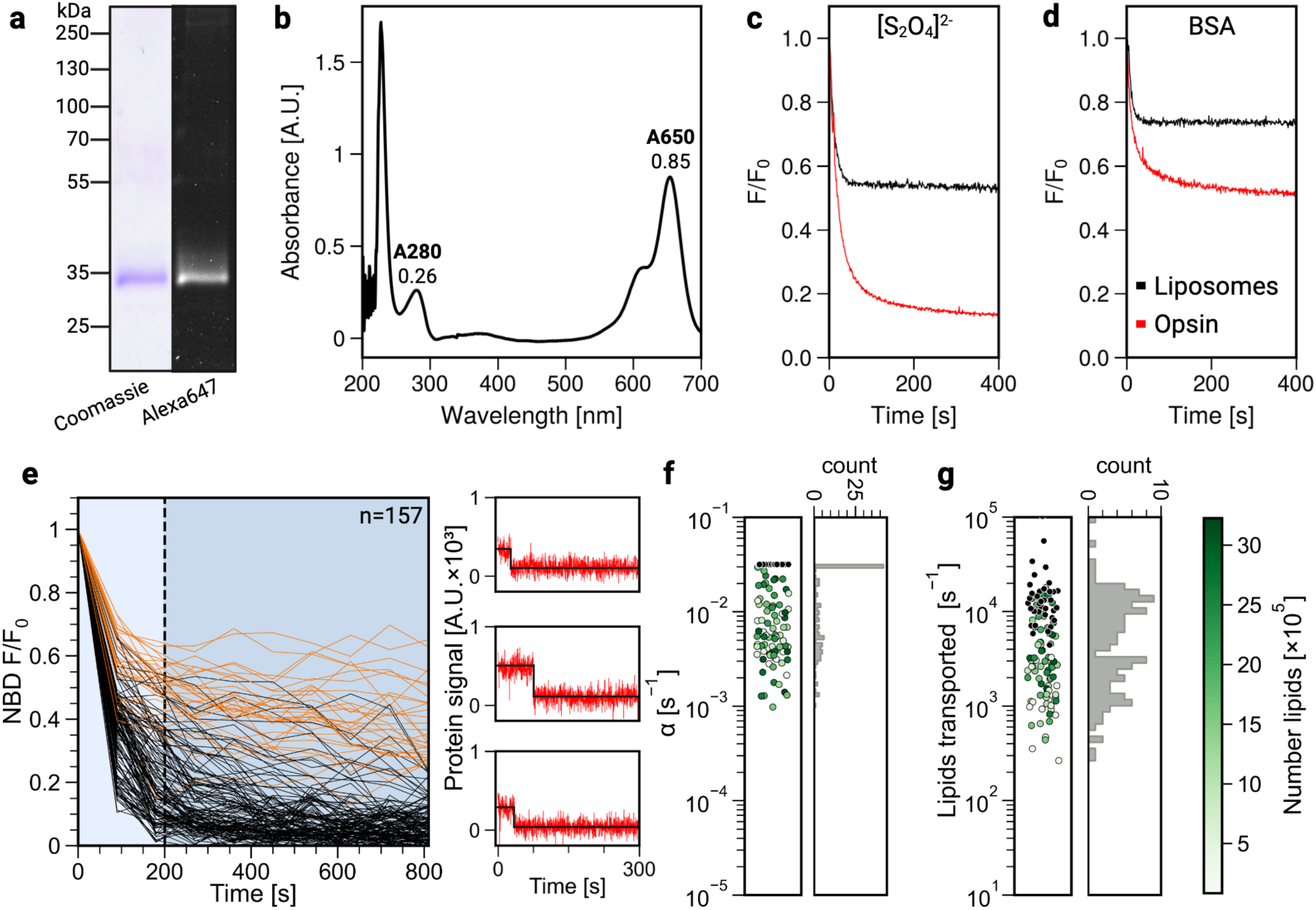
Single vesicle analysis of opsin-mediated scrambling. **a**, Opsin, purified from bovine retina, was labeled with Alexa Fluor 647 maleimide and analyzed by SDS-PAGE, visualized using Coomassie staining and in-gel fluorescence. **b**, Absorbance spectrum of Alexa Fluor 647-labeled opsin showing peaks corresponding to opsin (280 nm) and Alexa 647 (650 nm). Labeling stoichiometry of 1:1 was determined based on this spectrum. **c**,**d**, Alexa Fluor 647-labeled opsin was reconstituted into NBD-PC-containing LUVs at a protein/phospholipid ratio of 3.6 mg/mmol. Cuvette-based assays of opsin-mediated lipid scrambling probed using sodium dithionite ([S_2_O_4_]^2-^) (**c**) and BSA (**d**), added at t=0 s. Mock-reconstituted protein-free liposomes were analyzed alongside. **e**, Single vesicle scramblase assay of Alexa Fluor 647-labeled opsin proteoliposomes. Time-dependent decay of NBD fluorescence is shown for n=157 vesicles containing a single Alexa Fluor 647-labeled opsin, i.e., exhibiting single-step Alexa Fluor 647 photobleaching as seen in the three representative bleaching step analyses shown on the right. The majority of the vesicles (75%) retained <20% of the initial lipid signal at the end of the assay (black traces; remaining traces are shown in orange). **f**, High-activity traces (black lines in **e**) were fitted using a three-compartment model (Model 1) to obtain the scrambling rate constant α. Fitting revealed that a large number of traces reached the maximum α value possible within Model 1 (0.0317 s⁻¹). These are indicated by black symbols (left panel; the remaining symbols are color-coded to indicate number of lipids per vesicle according to the color map in panel (**g**)) and also reflected in the histogram of α values (right panel). **g**, Lipid transport rates per second for individual active vesicles containing a single opsin, color-coded according to the number of lipids per vesicle (color map on the right). Black symbols represent the lower limits to the scrambling rate as they correspond to traces where the α value reached the maximum allowed by the model.

**Extended Data Table 1.**
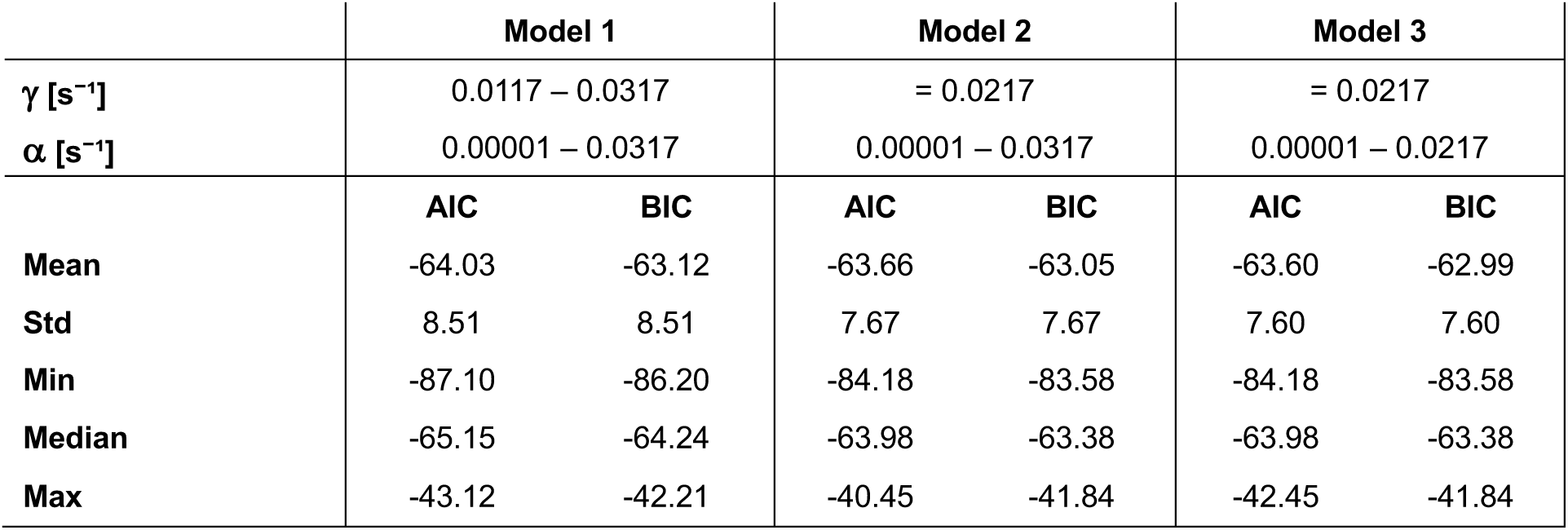
Comparison of models for fitting fluorescence decay traces. Three fitting models were tested. As the data are normalized, the models were constrained to the initial value of (t, F/F_0_)=(0,1). Also, the fluorescence plateau reached for protein-free liposomes was constrained to 0.40–0.60. The three models used different combinations of constraints for the extraction rate constant γ and the scrambling rate constant α. In Model 1, 0.0117 ≤ γ ≤ 0.0317 s⁻¹ was used to bracket the determined extraction rate (0.0217 ± 0.01 s⁻¹, Extended Data Fig. 5d). For Models 2 and 3, a fixed value of γ = 0.0217 s⁻¹ was used to assess potential overfitting effects of Model 1. Models 1–3 tested the effect of constraining α as follows: 0.00001 ≤ α ≤ 0.0317 s⁻¹ (Models 1, 2) and 0.00001 ≤ α ≤ 0.0217 s⁻¹ (Model 3), with 0.0317 s⁻¹ corresponding to the upper bound of our setup (i.e., highest γ value allowed) and 0.0217 s⁻¹ corresponding to the measured extraction rate. Akaike and Bayesian Information Criteria (AIC, BIC) were determined for every trace corresponding to vesicles with a single VDAC1 dimer (two Alexa Fluor 647 bleaching steps), and the statistics were computed. The lowest values were obtained for Model 1, which was then used for subsequent analyses.

|  | Model 1 |  | Model 2 |  | Model 3 |  |
| --- | --- | --- | --- | --- | --- | --- |
| $\gamma [\text{s}^{-1}]$ | 0.0117 – 0.0317 | | = 0.0217 | | = 0.0217 | |
| $\alpha [\text{s}^{-1}]$ | 0.00001 – 0.0317 | | 0.00001 – 0.0317 | | 0.00001 – 0.0217 | |
|  | AIC | BIC | AIC | BIC | AIC | BIC |
| Mean | -64.03 | -63.12 | -63.66 | -63.05 | -63.60 | -62.99 |
| Std | 8.51 | 8.51 | 7.67 | 7.67 | 7.60 | 7.60 |
| Min | -87.10 | -86.20 | -84.18 | -83.58 | -84.18 | -83.58 |
| Median | -65.15 | -64.24 | -63.98 | -63.38 | -63.98 | -63.38 |
| Max | -43.12 | -42.21 | -40.45 | -41.84 | -42.45 | -41.84 |

